# Partial reuse of circadian clock genes along parallel clines of diapause in two moth species

**DOI:** 10.1101/2022.06.22.497096

**Authors:** Yue Yu, Li-Li Huang, Fang-Sen Xue, Erik B. Dopman

## Abstract

Understanding the molecular basis of repeated evolution is essential for improving our ability to predict evolution. Genes repeatedly used in independent cases of adaptation to similar environments are strong candidates for predicting adaptation across phylogeny. The Asian corn borer (*Ostrinia furnacalis*; ACB) and the European corn borer (*Ostrinia nubilalis*; ECB) are two closely related moths that display remarkable adaptability to a wide range of climate on two separate continents, largely manifesting as changes in the timing of diapause (dormancy), but the genetic basis of parallel clinal responses remains to be characterized. We extensively sampled the ACB cline in China in a genome-wide association study (GWAS) using pooled sequencing data (Pool-seq). We characterized the genetic basis of clinal diapause response in ACB and showed that genes involved in circadian rhythm were over-represented among the candidate genes under spatially varying selection. Comparing with previous results from ECB, we found that the circadian clock gene period (*per*), but not pigment-dispersing factor receptor (*Pdfr*), was repeatedly used, but the alleles were not shared between the species. The corn borers’ shared adaptability is likely based in *per* but seemingly through independent mutational paths.

## INTRODUCTION

Elucidating the genetic mechanism of adaptive response to new environmental challenges is not only one of the fundamental goals in modern evolutionary biology, but also especially relevant in the face of accelerated climate change (Hoffmann & Sgrò, 2011; Sutherland et al., 2013; Williams et al., 2017). Phenotypic and genetic changes along a geographic transect (cline) are thought to be response to spatially varying selection. When such changes repeatedly occur, for example along another cline, it provides especially strong evidence that those reused targets of selection are at the core of the adaptability (Adrion et al., 2015; Arendt & Reznick, 2008; Machado et al., 2016; Stern & Orgogozo, 2009; Storz, 2016). Meta-analysis of the literature shows that the repeatability of evolution at the molecular level could be as high as 80% but decreases as the time since divergence increases (Conte et al., 2012; Storz, 2016). This suggests that closely related species might be more likely to arrive at a similar adaptive outcome due to the evolutionary history they more likely share – e.g., similar genetic background and constraints, access to ancestral polymorphisms, or a weaker reproductive barrier to exchanging beneficial alleles (Bohutínská et al., 2021; Lachapelle et al., 2015; Mallet, 2005) .

Adaptation to seasonality in insects often involves changes in diapause, a programmed developmental arrest. In climates where the distinct seasonal changes impose strong selection on insect phenology, the timing of diapause is critical to coping with *anticipated* adverse conditions and synchronizing the life cycle with local biotic factors such as the host plant (Anduaga et al., 2018; Beck, 1983; Bradshaw & Holzapfel, 2010; Denlinger et al., 2017; French et al., 2014; Koštál, 2006; D. C. Smith, 1988; M. J. Tauber et al., 1986). The consistent, predictable latitude-specific photoperiod is commonly the dominant cue for diapause, with temperature and humidity playing an important role in many cases. The latitudinal cline of diapause is well documented in many insects (e.g. Bean et al., 2012; Bradshaw & Holzapfel, 2010; Y.-S. Chen et al., 2013; Danilevskiĭ, 1965; Paolucci et al., 2013; Sadakiyo & Ishihara, 2011; Schmidt, Matzkin, et al., 2005; Shimizu & Kawasaki, 2001; Sota, 1994; Timer et al., 2010; Urbanski et al., 2012; X.-P. Wang et al., 2012). However, comparative studies on parallel clines of diapause and the underlying genetic polymorphisms have been scarce. The Asian tiger mosquito (*Aedes albopictus*) underwent rapid range expansion and climatic adaptation after being introduced to the United States recently, forming a cline in its egg diapause parallel to that in Japan where it is native (Batz et al., 2020; Urbanski et al., 2012). Comparative genomics has just begun to identify the underlying genetic variations (Boyle et al., 2021). The model organism *Drosophila melanogaster* so far has the best characterized clines and candidate genes. Three temperate clines of ovarian diapause in *D. melanogaster* have been described in North America (Schmidt et al., 2005), Australia (S. F. Lee et al., 2011; see also Nagy et al., 2018 and Zonato et al., 2017), and Europe, with the European one being relatively shallow (Pegoraro et al., 2017). The gene couch potato (*cpo*) was found to underlie the diapause cline in North America (Cogni et al., 2014; Schmidt et al., 2008). In Australia, *cpo* was also clinal, although its association with the phenotype was not evident (Lee et al., 2011). On the other hand, a diapause-enhancing allele in timeless (*tim*) that recently spread from Italy might be the reason for the dampened cline in Europe (Pegoraro et al., 2017; Sandrelli et al., 2007; E. Tauber et al., 2007). Importantly, no single gene emerged as globally responsible for adaptive diapause variation. In the recent decade, affordable whole-genome sequencing of *Drosophila* and other species enabled large-scale association studies, which uncovered a plethora of natural variations under spatially varying selection (e.g. E. J. Dowle et al., 2020; Erickson et al., 2020; Fabian et al., 2012; Machado et al., 2016; Ragland et al., 2017; Reinhardt et al., 2014). This allows for genome-wide comparisons of clines to thoroughly test the repeatability of adaptation across geographic space. The North American and Australian *Drosophila* geographic clines overlapped by as much as 31% of clinal genes (Fabian et al., 2012) and 1.1% of all shared single nucleotide polymorphisms (SNP) (Reinhardt et al., 2014). Whether or how these genetic variations are connected to diapause remains to be seen, but many of these genes are from pathways implicated in the diapause program by either direct (gene knockdown) or indirect (phenotype-genotype association) analyses, such as the circadian clock pathway (including *tim*), the insulin signaling pathway, and the ecdysone signaling pathway (Denlinger, 2022; Fabian et al., 2012).

In this study, we take advantage of parallel clines of phenological adaptation in the Asian corn borer *Ostrinia furnacalis* (Guenée ) (Lepidoptera: Crambidae:) and the European corn borer *Ostrinia nubilalis* (Hübner) to elucidate the genetic basis of repeated evolution. The two closely related moths colonized maize after its introduction into Europe and Asia around 500 years ago, (S. Chen & Kung, 2016; Tenaillon & Charcosset, 2011). The Asian corn borer (ACB) is now found throughout Asia and Australia, and the European corn borer (ECB) in the U.S. and southern Canada in addition to Europe, North Africa, and western Asia where it is native. In their range of distribution across the temperate and even tropical climates, the number of generations per year (voltinism) increases with decreasing latitude, largely constrained by the length of the growing season. Towards the end of the growing season, before the onset of winter, both species enter diapause as a fifth instar larva and then resume development in the next spring or summer. Corn borer populations across latitude differ in the timing of both the start and end of diapause (Huang et al., 2020; Levy et al., 2015; Ma et al., 2008; McLeod, 1976; Showers et al., 1975), respectively measured by critical daylength (CDL), the photoperiod that induces diapause in half the population, and post-diapause development (PDD) time, the average number of days required to terminate diapause and pupate upon receiving summer-like cues. For both species, when raised in common garden conditions (constant photoperiod and temperature), CDL increases with increasing latitude of origin (Figure 1). On the other hand, PDD of both species varies with latitude non-monotonically, a pattern interpreted as reflecting the requirement of fitting discrete generations into the latitudinally-varying growing season length (the “sawtooth” pattern, Roff, 1980, also see Figure 1b in Levy et al., 2015).

**Figure 1.**
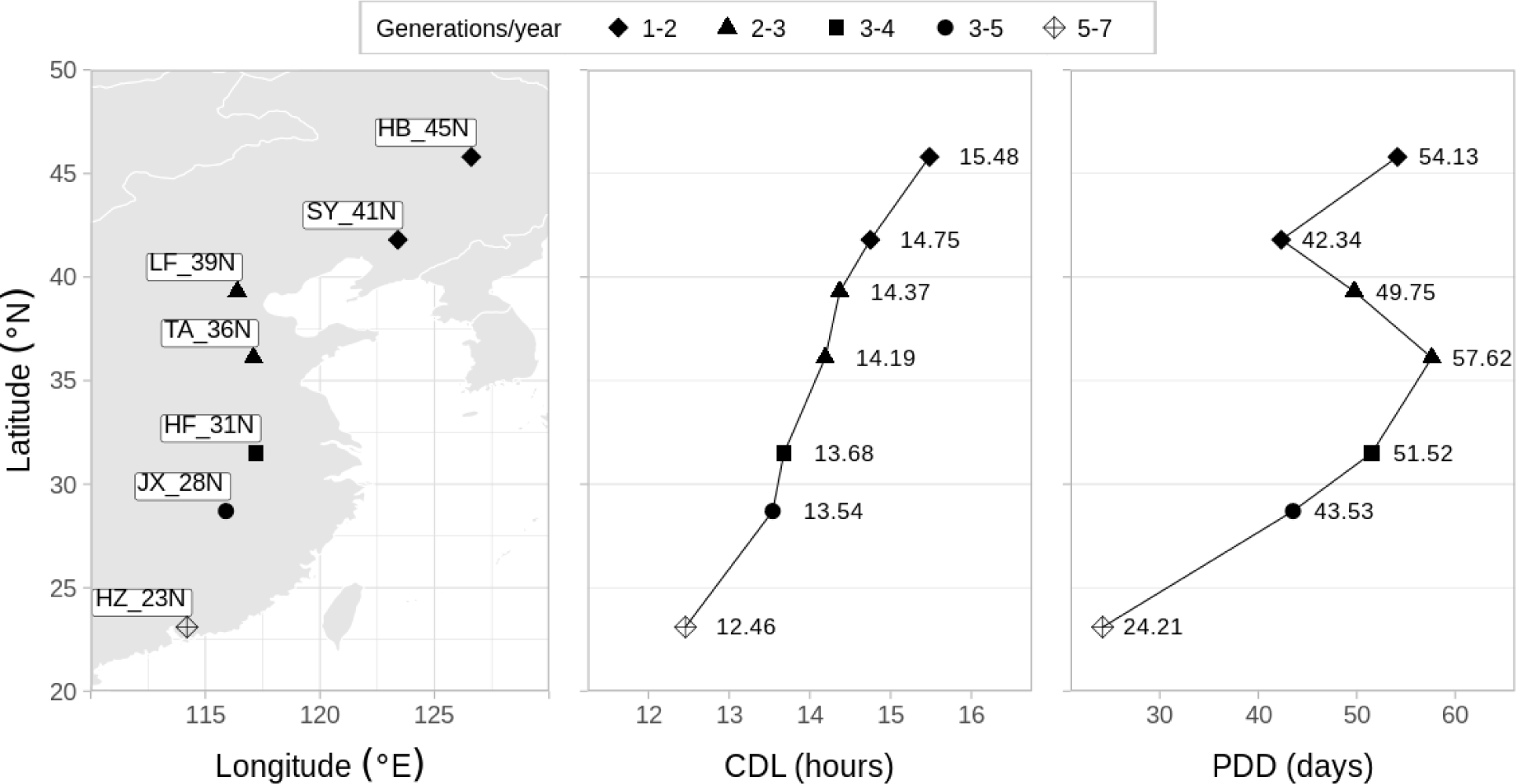
The location and population attributes of the sampled ACB populations. Population name abbreviations: HB_45N (Haerbin, 45.8°N, 126.6°E); SY_41N (Shenyang, 41.8°N, 123.4°E); LF_39N (Langfang, 39.3°N, 116.4°E); TA_36N (Tai’an, 36.1°N, 117.1°E); HF_31N (Hefei, 31.5°N, 117.2°E); JX_28N (Jiangxi, 28.7°N, 115.9°E); HZ_23N (Huizhou, 23.1°N, 114.2°E). CDL: critical daylength measured at 28°C. PDD: post -diapause development time measured under natural conditions during the winter of 2015 in Nanchang, Jiangxi, China.

More is known about genes underlying diapause in ECB than ACB. Population genetics revealed that the circadian clock genes period (*per*) and cryptochrome1 (*cry1*) exhibit sawtooth oscillations in allele frequency along a North American latitudinal transect coincident with transitions in voltinism (Levy et al., 2015). A more recent study identified two epistatically interacting quantitative trait loci (QTL) underlying PDD variation in ECB, each encompassing a circadian clock gene: the clinally varying gene *per* and Pigment-dispersing factor receptor (*Pdfr*), the receptor of the main clock cell neurotransmitter. Alleles of the two genes are highly correlated in natural populations (high linkage disequilibrium), despite being ∼4.5 Mb apart on the Z (sex) chromosome (Kozak et al., 2019). CDL variation was mapped to a polymorphic chromosomal inversion containing one of the two PDD QTLs (Golczer, 2019) and was further placed near *per* using collinear strains (Yu & Dopman, in preparation). In contrast, candidate diapause genes in ACB have yet to be identified.

In addition to whether there is repeated genetic evolution in these corn borer lineages, a critical question is how the ability to rapidly adapt to the wide range of climates arose. Standing genetic variations are thought to facilitate rapid evolutionary response as they are immediately available (unlike new beneficial mutations) and potentially have been “tested” in past selective events (Barrett & Schluter, 2008). As a classic example, the threespine stickleback repeatedly adapted to freshwater environments, evolving a reduced number of lateral plates in parallel based on the same ancestral allele from the marine ancestor (Colosimo et al., 2005). Introgression is another way for a species to utilize preexisting variations (Mallet, 2005). In the fly *Rhagoletis pomonella*, a chromosomal inversion introgressed from Mexico to the U.S. seemed to have facilitated clinal adaptation and host plant shifts which involved changes in diapause timing (Feder et al., 2003; Michel et al., 2007). In the corn borers, there is evidence of hybridization and introgression in areas of sympatry between ACB and ECB in the Yili region, China (Y. Wang et al., 2017) and between ACB and *Ostrinia scapulalis* (sibling species to ECB) in Zhangjiakou, China (Bourguet et al., 2014; Frolov et al., 2007; W.-D. Li et al., 2003). It is unclear how much of a role adaptive introgression plays in parallel phenotypic evolution of ACB and ECB.

Here, we use 7 ACB populations sampled across 2,764 kilometers and 23 degrees of latitude to investigate three questions: (i) What is the genetic architecture of the diapause response in ACB? (ii) Is there a repeated use of genes underlying the parallel phenotypic evolution of ACB and ECB? (iii) How did the genetic polymorphisms underlying the parallel evolution arise? We hypothesize that *per* and *Pdfr* underlie the latitudinal variation of diapause in ACB and the genetic variations might have introgressed between ACB and ECB. With pooled whole-genome sequencing data, we describe the population structure and highlight some strong candidate genes for the diapause variation. We quantify the level of introgression between ACB and ECB and identify an introgressed locus that might have been adaptive.

## MATERIALS AND METHODS

### Pool-seq data across latitudes

#### ACB sampling and phenotyping

Diapausing ACB larvae were collected from 7 populations in maize fields along the latitudinal transect in China in October, 2015, spaced 3.78 degrees of latitude (503.96 kilometers) apart on average (Figure 1). The populations are named according to their place of origin and latitude, herein abbreviated as: HB_45N (Haerbin, 45.8°N, 126.6°E); SY_41N (Shenyang, 41.8°N, 123.4°E); LF_39N (Langfang, 39.3°N, 116.4°E); TA_36N (Tai ’an, 36.1°N, 117.1°E); HF_31N (Hefei, 31.5°N, 117.2°E); JX_28N (Jiangxi, 28.7°N, 115.9°E); HZ_23N (Huizhou, 23.1°N, 114.2°E). In a common garden experiment, the collected diapausing larvae were exposed to and kept under the natural conditions in Nanchang, Jiangxi Province, and the average number of days to pupation after exposure (PDD) was recorded for each population. Their offspring were reared under various constant photoperiods (LD 11:13 to 18:6) at 28°C and the critical daylength (CDL) was determined from the photoperiodic response curve of diapause incidence (Huang et al., 2020).

Adults were preserved in AllTissue Protect and DNA was extracted from their lower abdomens using the Qiagen DNeasy Blood & Tissue Kit and quantified using Qubit (Thermo-Fisher). For each population, equimolar amounts of DNA from 30 males (the homogametic sex) were pooled together for sequencing. Although a pool of at least 40 individuals was considered to be best for reducing sampling bias (Schlötterer et al., 2014) , studies found a pool size of 30 to be sufficient (Boitard et al., 2012; Gautier et al., 2013).

#### Sequencing data and alignment

The 7 libraries were prepared for paired-end 150 bp sequencing on Illumina HiSeq 4000 at Novogene (Beijing, China). We used Trimmomatic (Bolger et al., 2014) to trim the reads for Illumina adapter sequences and quality (*LEADING:3 TRAILING:3 SLIDINGWINDOW:4:15 MINLEN:36*). Raw and trimmed data quality was checked using FastQC (Andrews, 2010). Trimmed reads were aligned to the 437.3 Mb representative ACB genome (assembly name ASM419383v1, RefSeq accession GCF_004193835.1; soft-masked) using Bowtie2 (Langmead & Salzberg, 2012) with the very-sensitive end-to-end alignment setting. Duplicate reads were removed using the Picard MarkDuplicates tool (*Picard Toolkit*, 2019). Samtools (H. Li et al., 2009) was used to filter for mapping quality (*-q 20 -F 0x0004*) and merge the individual bam files into the ‘mpileup’ format. We used Popoolation2 (Kofler et al., 2011) to identify and filter out indels.

To arrange the ACB reference scaffolds in chromosomes for visualizing genome-wide statistics later, we aligned the ACB scaffolds to the silkworm (*Bombyx mori*) chromosomes. *B. mori* is the best studied Lepidopteran genome and Lepidopteran genomes are largely syntenic, except that the *B. mori* Chromosomes 11, 23, and 24 are the result of 3 fusion events of 6 ancestral Lepidopteran chromosomes (Yasukochi et al., 2016). We first identified pairs of ACB gene models (RefSeq GCF_004193835.1 Protein FASTA) and silkworm gene models from Silkbase (version Jan 2017; Kawamoto et al., 2019) that were reciprocal best BLAST hits of each other (Altschul et al., 1990). The order of the aligned genes was used to orient the scaffolds against the *B. mori* chromosomes. Of the 7,721 ACB nuclear scaffolds, 6,070 had no predicted genes; 534 contained predicted genes and unambiguously mapped to a chromosome; for the scaffolds with genes that did not all align to only one chromosome, we assigned the scaffold to the chromosome with the longest alignment, resulting in an additional 644 mapped scaffolds. These 1,178 mapped scaffolds were on average 351 kb long, adding up to 95% of the length of the ACB reference genome. In contrast, unplaced scaffolds averaged 7.5 kb in length (SNP-containing only, n = 2,552).

Prior to any analysis, the mitochondrial scaffold (NC_003368.1), which was retained only for read alignment, was filtered out from the data.

### Genetic basis of local adaptation

Only biallelic SNPs were used for all statistics mentioned below except *d_xy_*. To assess population structure and compare with the *XtX* statistic (see below), *F*_ST_ was calculated using Popoolation2 (Kofler et al., 2011) for single SNPs covered by between 15 and 200 reads in every population as well as for the genome-wide average (see Supplemental File S1). We defined *F*_ST_ outliers as those in the top 0.01% of the empirical distribution for at least three of the population pairs (Cavedon et al., 2019).

We used BayPass 2.2 (Gautier, 2015) to detect signatures of selection while controlling for the unknown shared demographic history among the populations, which is inferred from the covariance matrix of allele frequencies, Ω (Gautier, 2015; Günther & Coop, 2013) . The statistic *XtX* measures allele frequency differentiation across all populations after accounting for Ω (Gautier, 2015; Günther & Coop, 2013) . First, we converted Popoolation’s output *sync* file to the *genobaypass* file format required by BayPass using the R (R Core Team, 2020) function poolfstat::popsync2pooldata with the settings *min.cov.per.pool=3* and *min.maf=0.05*, which filtered the SNPs by read coverage and minor allele frequency, respectively. Then, *XtX* statistic was calculated for each SNP under the core model in BayPass. We made a pseudo-observed dataset (POD) of 100,000 SNPs with BayPass’s R function simulate.baypass (*coverage=26*) to sample new SNPs under the model parameters estimated from the full data. *XtX* under the core model was calculated for the simulated neutral SNPs in the POD. The 99.99% percentile (*XtX* = 26.194) of this posterior predictive distribution of *XtX* was used as the significance threshold for the full data. Like *F*_ST_, low *XtX* outliers can be used to detect balancing selection but there were no SNPs in the full data below the POD’s lower 0.01% percentile (*XtX* = 1.428).

The covariance matrix Ω estimated by BayPass above was used to visualize the inferred demographic history (with no gene flow) by first converting it to a correlation matrix in R with the function cov2cor, then to a distance matrix by calculating d_ij_ = 1 – |ρ_ij_|, where d_ij_ are elements of the distance matrix and ρ_ij_ are the corresponding elements in the correlation matrix (Gautier, 2015). And finally, this distance matrix was used for UPGMA clustering by the R function hclust to construct a phylogenetic tree without gene flow.

For the genetic basis underlying each variable of interest, we used BayPass to identify SNPs associated with four population covariates while accounting for the shared population history. The four covariates were latitude, average number of generations per year (voltinism), CDL measured at 28^°^C, and average PDD measured under natural conditions (Figure 1). We tested for pairwise correlation between the covariates and found high correlations (|Spearman’s rho| <0.429 between PDD and the other covariates; |rho| > 0.981 for all other pairs; Supplemental Table S1). Therefore, we ran the Importance Sampling (IS) covariate model which was recommended for cases with inter-correlated covariates and < 8 populations (Gautier, 2019). This model is equivalent to several separate univariate regression models. We specified the options *d_0_y_ij_=12* (a fifth of the haploid pool size) and *scalecov* (to standardize the covariates). We used two cut-offs for significant association: Bayes Factor (BF) > 30 deciban units (dB), when BF > 20 dB is considered “decisive” evidence for there existing a linear SNP-covariate association rather than not (Jeffreys’ rule; Jeffreys, 1998), and the empirical Bayesian probability eBPis > 3, which is roughly equivalent to a probability < 0.001% that the regression coefficient β_i_ is 0 in the frequentist view (Gautier, 2015).

**Table 1.** Number of SNPs significantly associated with covariates.

| Covariate | Total SNPs<br>(in predicted genes) | SNPs in CDS | XtX outliers<br>(in predicted genes) |
| --- | --- | --- | --- |
| Latitude | 633 (126) | 171 | 117 (9) |
| Average voltinism | 813 (167) | 225 | 134 (18) |
| CDL | 592 (122) | 169 | 115 (11) |
| PDD | 474 (149) | 155 | 82 (22) |

### Signatures of introgression

We used four-taxon tests to investigate adaptive introgression between ACB and ECB. We calculated the statistics *D* (Durand et al., 2011; Green et al., 2010) and *f_d_* (Martin et al., 2015) using a custom R script following equations in Martin et al. (2015). Briefly, given an evolutionary relationship of 3 populations and an outgroup, (((P_1_, P_2_), P_3_), O), the ABBA BABA test (the *D*-statistic) measures introgression between P_3_ and the ingroups P_1_/P_2_ by comparing the number of sites with the pattern ABBA (P_2_ and P_3_ sharing the derived allele “B”) with the number of BABA sites (P_1_ and P_3_ sharing the derived allele “B”). In the absence of introgression, the ABBA and BABA patterns are expected to occur equally frequently under incomplete lineage sorting. An excess of shared derived alleles indicates introgression between P_2_ and P_3_ (ABBA > BABA; *D* > 0) or between P_1_ and P_3_ (ABBA < BABA; *D* < 0). In the case of population genomics data, relative frequencies of the patterns are calculated from allele frequencies (Durand et al., 2011; Green et al., 2010; Martin et al., 2015). *D* does not perform well in small genomic windows due to its sensitivity to the local nucleotide diversity, therefore we also used the *f_d_* statistic for small genomic windows (Martin et al., 2015). The *f_d_* statistic measures the proportion of introgression between P_2_ and P_3_ (i.e., in windows where *D* > 0) compared to the scenario of complete homogenization of allele frequencies between P_2_ and P_3_.

For our four-taxon tests, we used our ACB populations and two populations of the ECB as the ingroups and the American Lotus Borer (*Ostrinia penitalis*; herein ALB) as the outgroup.

Publicly available pool-seq data of the ECB populations EA (East Aurora, NY, USA; herein ECB_EA; SRA# SRX5775029) and PY (Penn Yan, NY, USA; herein ECB_PY; SRA# SRX5775027) (Kozak et al., 2019) and 3 individually sequenced whole-genome data of ALB (SRA# SRX8977671, SRX8977672, SRX8977673) were downloaded from the NCBI SRA database. We aligned all genomic data to the ACB RefSeq genome using the same pipeline described above, except that for ALB we used the very-sensitive local alignment setting in Bowtie2. We only retained SNPs that were fixed in the 3 ALB individuals to infer the ancestral states.

We first calculated genome-wide average *D* to look at the overall extent of introgression and to help select tree topologies meaningful for the calculation of *f_d_* later. We tested all possible four-taxon trees that were of the structures (((ECB, ECB), ACB), ALB) and (((ACB, ACB), ECB), ALB) where ECB was either ECB_EA or ECB_PY and ACB was any of the 7 ACB populations. We performed block jackknifing in blocks of 6,000 SNPs to obtain the standard error and *z*-score for the *D* values. We considered a *D* value to be significantly different from 0 if *z* ≥ 3.

To identify regions of elevated introgression relative to the rest of the genome that might be related to diapause, we calculated *f_d_* for the 6 possible tree topologies of the structure (((HZ_23N, ACB), ECB_PY), ALB), where ACB (P_2_) was any of the non-HZ_23N populations. We decided to use HZ_23N as P_1_ and ECB_PY as P_3_ because (i) genome-wide *D* from trees with ECB as the ingroups showed that there had been more of the genome introgressed between ECB_PY and ACB than between ECB_EA and ACB, (ii) genome-wide *D* from trees with ACB as the ingroups showed that HZ_23N, relative to the other ACB populations, introgressed the least with ECB, (iii) *f_d_* is only meaningful when *D* > 0 (more P_2_-P_3_ introgression than P_1_-P_3_) (Martin et al., 2015), and (iv) HZ_23N was phenotypically the closest to nondiapausing and genetically the most differentiated from the other populations. We calculated *f_d_* in non-overlapping windows of 300 SNPs (median length = 22,714 bp) to avoid stochastic errors that might occur in windows with too few SNPs (Martin, 2018). We defined *f_d_* outliers as the top 1% of the empirical distribution after filtering for the informative windows (*D* ≥ 0, number of SNPs > 150), and *f_d_* outliers that additionally showed signatures of selection (containing *XtX* outliers) were considered to show evidence of adaptive introgression.

The absolute measure of divergence, *d_xy_*, was used to differentiate between introgression and shared ancient polymorphisms (Martin et al., 2015; J. Smith & Kronforst, 2013), as both can result in an excess of shared derived alleles (Eriksson & Manica, 2012). We estimated average *d_xy_* from monomorphic and biallelic sites between pairs of ACB and ECB_PY in non- overlapping windows of 25000 bp using a custom R script, following the equation based on allele frequencies in J. Smith & Kronforst (2013). Higher *d_xy_* than the rest of the genome was expected for the scenario of shared ancient polymorphism.

### Gene annotation

To functionally annotate the SNPs or genomic windows of interest from the analyses described above, we first used BEDTools (Quinlan & Hall, 2010) to map them to the genome annotations (NCBI *Ostrinia furnacalis* Annotation Release 100). Specifically, we used *bedtools window* to annotate SNPs/windows if they were within 1,000 bp of the genes in the *gff*3 annotation file.

Note that the majority of the genes were predicted by Gnomon (Sourvorov et al., 2010) through the NCBI Eukaryotic Genome Annotation Pipeline (herein, “predicted ACB genes”).

Besides the Gnomon annotations, we associated Gene Ontology (GO) terms to the genes using PANTHER (Mi et al., 2019). The predicted ACB protein sequences were scored against gene models in the PANTHER HMM Library 16.0 with an HMM E-value cut-off of 1e-23, which is considered high confidence (Mi et al., 2019). The resulting PANTHER Generic Mapping files were uploaded to the PANTHER website to obtain the list of PANTHER protein families, any members of these families that are from the fruit fly (*Drosophila melanogaster*), and the associated GO terms. To complement the PANTHER annotations, we also performed BLAST against the *D. melanogaster* genome in the FlyBase database (release FB2021_02; *D. melanogaster* r6.39) (Larkin et al., 2021) using a conventional E-value cut-off of 1e-50.

To identify significantly enriched GO terms and pathways, we uploaded the *D. melanogaster* gene symbols obtained above to g:Profiler (Raudvere et al., 2019) as “multiquery”, selected “only annotated genes” from *D. melanogaster* as the background, Benjamini-Hochberg FDR-adjusted p < 0.05 for significance, GO Biological Process and KEGG as the sources of annotation, and limited the list to term size 10-500. We also used GeneCodis4 (Garcia-Moreno et al., 2022) with the Reactome pathway database as the source of annotation, which was unavailable for *D. melanogaster* in g:Profiler. The lists of significantly enriched GO-BP terms from g:Profiler (*p_adj_* < 0.05) were summarized in REVIGO through clustering in a two-dimensional semantic space with a dispensability cut-off of 0.5 (Supek et al., 2011).

Dispensability is based on the pairwise similarity score SimRel (Schlicker et al., 2006) and determines which terms are assigned to the same cluster. REVIGO selects the representative GO term for a cluster based on low dispensability, low frequency of occurrence in the database, high enrichment, and any parental relationship to other GO terms in the same cluster (Supek et al., 2011).

### R packages

We used the following `R` (R Core Team, 2020) packages: `ade4` (Chessel et al., 2004) for the Mantel test and `poolfstat` (Hivert et al., 2018) for BayPass-related analyses; `data.table` (M. Dowle & Srinivasan, 2020), `dplyr` (Wickham et al., 2020), `forcats` (Ammar, 2020), and `tidyr` (Wickham, 2020) for data manipulation; `ape` (Paradis & Schliep, 2019), `cowplot` (Wilke, 2020), `ggplot2` (Wickham, 2016), `ggpubr` (Kassambara, 2020), `ggrepel` (Slowikowski, 2021), `ggtree` (Yu, 2020), `maps` (Brownrigg et al., 2018), `randomcoloR` (Ammar, 2019), and `VennDiagram` (H. Chen & Boutros, 2011) for visualization.

## RESULTS

### Sequence alignment

To identify genomic regions potentially underlying local adaptation and diapause traits, we tested for genome-wide association with population-specific attributes using pooled sequencing data from 7 ACB populations across latitudes (n = 30 males per pool; Figure 1). Paired-end 150 bp Illumina reads were aligned to the ACB reference genome (RefSeq accession GCF_004193835.1). We obtained an average breadth of coverage of 88.5% (ranging from 87.5% to 89.4% across the 7 pools); the average depth of coverage was 26.5 reads per covered position (ranging from 24.3X to 30.0X across the 7 pools).

### Population structure

We characterized the population structure in the 7 ACB populations using *F*_ST_ as well as the Bayesian estimation of a covariance matrix and found that the population structure could not be explained by isolation-by-distance alone. Based on a total of 6,584,518 biallelic SNPs, the genome-wide average *F*_ST_ between populations ranged from 0.048 to 0.074 (Figure 2A). When F_ST_/(1- F_ST_) was plotted against pairwise geographic distance (Rousset, 1997), isolation-by-distance was not statistically significant at 0.05 (Mantel test with 10,000 replications, *r* = 0.437, *p* = 0.064) (Figure 2C). Population structure captured in the covariance matrix of population allele frequencies (Ω, estimated by BayPass) further supported a complex relationship among populations, in which populations did not seem to cluster by geographic proximity (Figure 2B&D). For example, the second northernmost (SY_41N) and second southernmost (JX_28N) populations clustered together.

**Figure 2.**
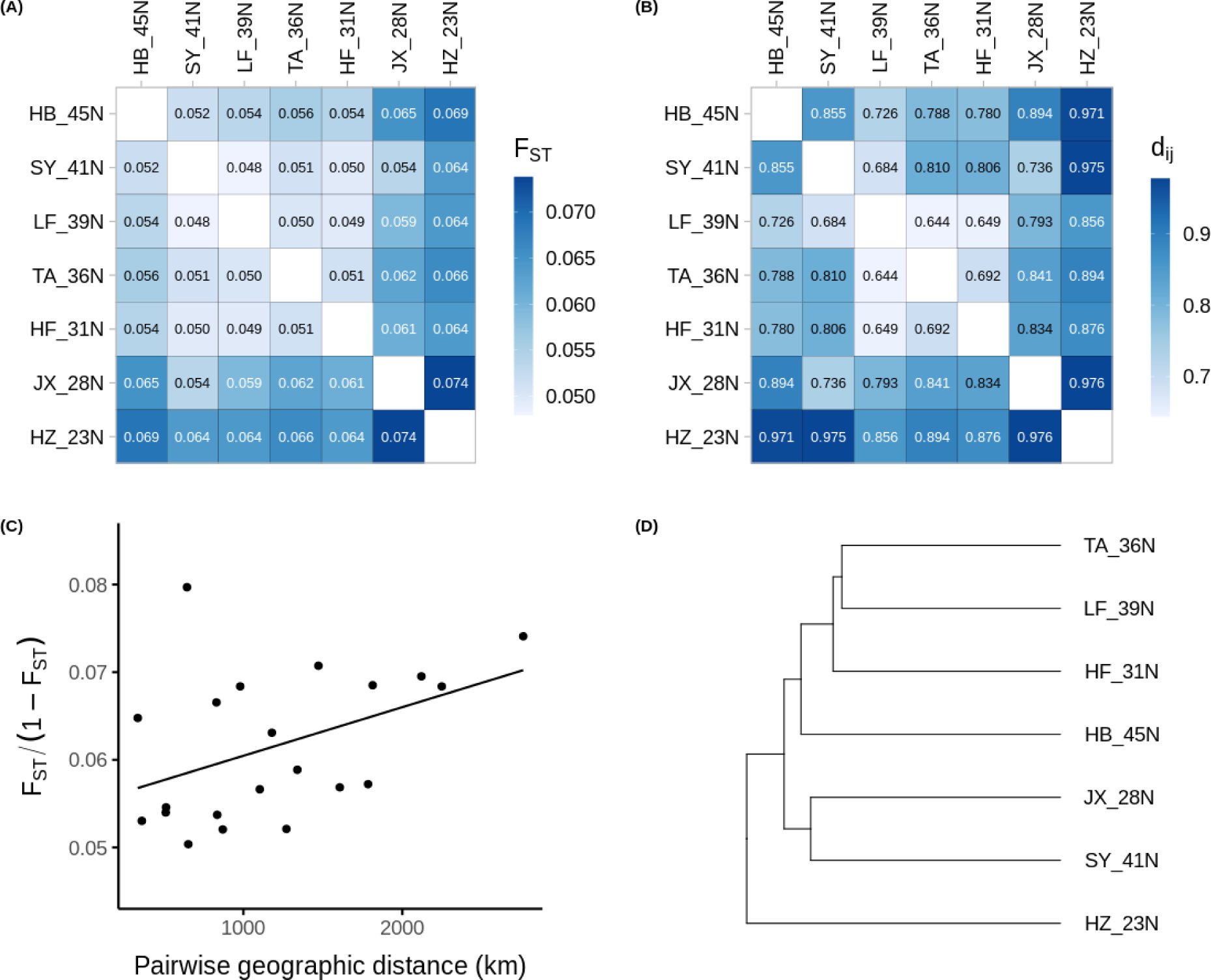
(A) Pairwise *F*_ST_ among the 7 ACB populations. (B) Distance matrix (d_ij_) converted from Ω, the scaled covariance matrix estimated by BayPass. (C) *F*_ST_ /(1 – *F*_ST_) plotted against the geographic distance between pairs of populations (Rousset, 1997) to test isolation-by-distance (Mantel test with 10,000 replications, *r* = 0.437, *p* = 0.064). (D) Estimated demographic history (without gene flow) using UPGMA hierarchical clustering based on (B).

Of the four covariates (latitude, voltinism, CDL, and PDD), only PDD (non-significantly) correlated with the population structure, namely the first Principal Component of Ω (Spearman’s rho = -0.75, *p* = 0.052; |rho| < 0.1 for the other covariates; Supplemental Table S1). The implication is that there could potentially be an increased false positive rate in SNP-PDD associations (Frachon et al., 2018; Günther & Coop, 2013) .

### Signature of selection

We used BayPass to calculate *XtX*, which is a statistic similar to *F*_ST_, measuring a SNP’s differentiation among all populations while controlling for population structure (Ω). Out of a total of 6,758,085 SNPs considered by BayPass, there were 282 *XtX* outliers (top 0.01% POD threshold, *XtX* > 26.194). Comparing with the widely-used method of pairwise *F*_ST_ outlier test, 161 of the 282 *XtX* outliers were also *F*_ST_ outliers, which we defined as SNPs in the top 0.01% of the *F*_ST_ distribution for more than 2 pairs of populations (1,119 *F*_ST_ outliers in total). The *XtX* outliers were found on a majority of the chromosomes but were most dense in the Z chromosome (Supplemental Figure S1).

Of the 282 *XtX* outliers, 183 were in or close to (within 1,000 bp) a predicted gene (Supplemental File S3); furthermore, 119 of them were within predicted protein coding regions, in contrast to 9.4% of the genome predicted to be CDS (NCBI *Ostrinia furnacalis* Annotation Release 100). For the 67 predicted ACB genes that the *XtX* outliers were mapped to, 39 had an ortholog in *D. melanogaster* (based on joint consideration of PANTHER and BLAST results). Semantic clustering of the significantly enriched GO terms (*p_adj_* < 0.05) by REVIGO identified representative GO terms from biological processes involving the circadian clock (GO:0042220, GO:0048511, GO:0032922), sensory system development (GO:0007423, GO:0048880), and metabolic processes (GO:0046434, GO:0031327) (Supplemental File S4). Notably, the circadian clock pathway was significantly over-represented, with 4 genes annotated in the Circadian Clock pathway in Reactome (R-DME-432626, adjusted *p* = 0.038): *Clk*, *per*, *wdb*, and *Rpt2* (also known as 26S proteasome regulatory subunit 4, or *Pros26.4*); *Clk* and *per* were also annotated in the Circadian Rhythm pathway in KEGG (KEGG:04711, *p_adj_* = 0.049).

### Significant association with phenology

We identified SNPs whose allele frequencies significantly covaried with the geographic and phenotypic covariates under the IS covariate model in BayPass using the thresholds BF > 30 dB and eBPis > 3 (Table 1; Figure 3). Under this model, the covariates were treated separately because they were correlated with each other to varying degrees (|Spearman’s rho| < 0.429 between PDD and the other covariates; |rho| > 0.981 for all other pairs; Supplemental Table S1). We focused on genes associated with CDL and PDD and looked for any additional genes that might be uniquely associated with voltinism or latitude.

**Figure 3.**
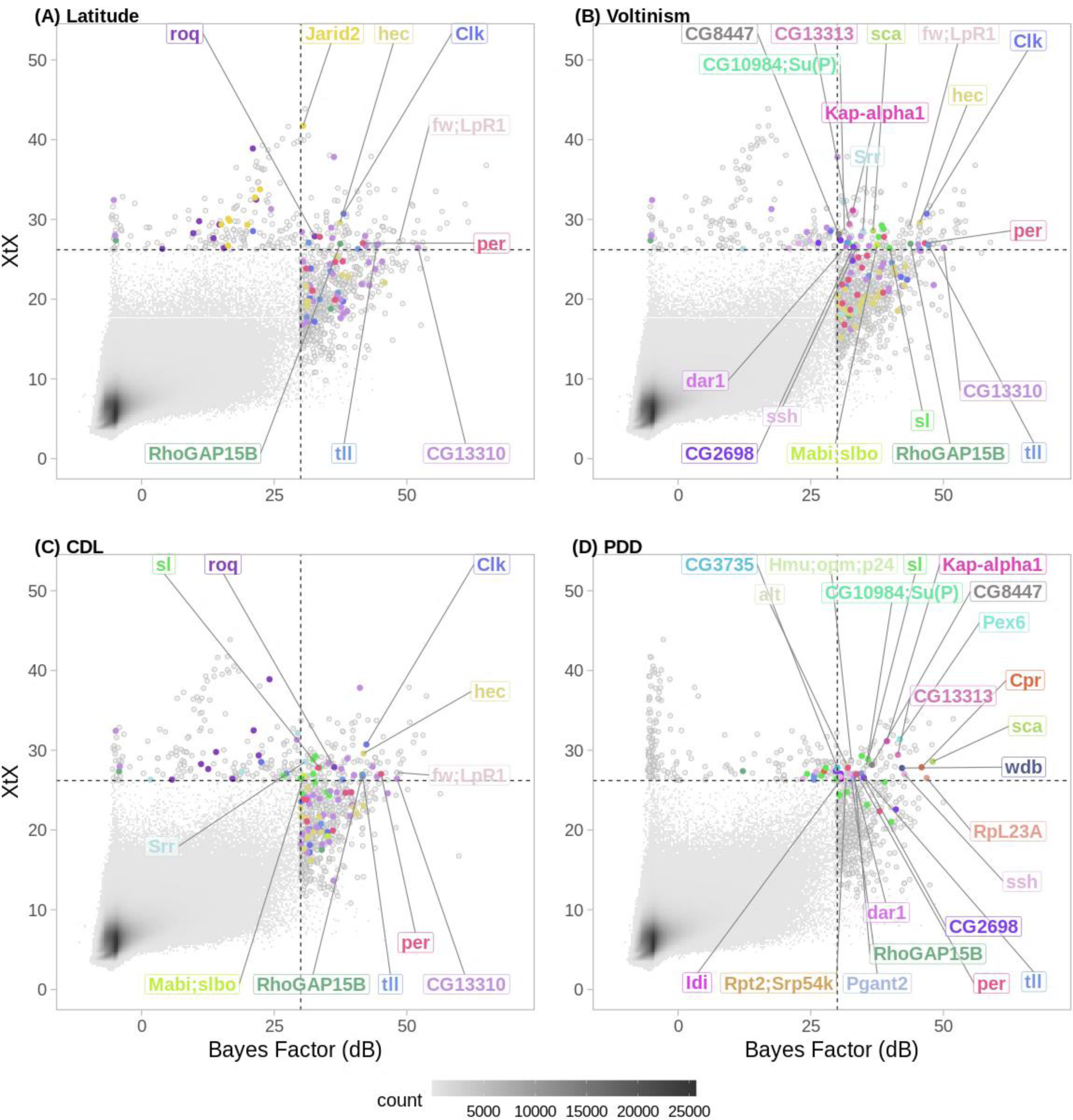
SNP-specific *XtX* (population differentiation) plotted against the Bayes Factor (SNP-covariate association) for each population covariate: (A) latitude of sampling location, (B) average number of generations per year (voltinism), (C) critical daylength (CDL) measured at 28°C, (D) post -diapause development (PDD) time measured under natural conditions in a common garden experiment. Dashed lines show significance thresholds (for *XtX*: 26.194; for covariates: BF > 30dB, and additionally eBPis > 3 which is not plotted on an axis). The large number of nonsignificant SNPs are plotted as a 2D histogram where the greyscale color of each grid represents the count of SNPs within the bin; the majority of these are in the lower left quadrant formed by threshold lines. SNPs significant for differentiation and/or association with a covariate are plotted individually (grey open circles or colored closed circles). Genes that contained a SNP significant for both (upper right quadrant, colored closed circles) are labeled; additional SNPs from these genes in the other two quadrants of significance (upper left, lower right) are also colored accordingly. Gene symbols separated by a semicolon indicate a SNP near more than one predicted ACB gene (within 1,000 bp) or the predicted ACB gene had more than one putative *D. melanogaster* homolog in the same PANTHER protein model.

#### CDL

In total, we found 592 SNPs significantly associated with CDL, mapped to 122 predicted genes (Table 1; Supplemental File S3). Among the significantly enriched GO terms, the least dispensable representative GO terms (REVIGO dispensability = 0) included circadian rhythm and its regulation (GO:0007623, GO:0042752), “inter-male aggressive behavior” (GO:0002121), “post-embryonic appendage morphogenesis” (GO:0035120), and actin filament-based processes (GO:0030029, GO:0030050) (see Supplemental File S4 for the full list). There were 115 CDL- associated SNPs that were also *XtX* outliers, mapped to 11 genes, including the aforementioned clock genes *Clk* and *per* (Figure 3C).

#### PDD

In total, we found 474 SNPs significantly associated with PDD, mapped to 149 predicted genes (Table 1; Supplemental File S3). The significantly enriched GO terms were represented by “calcium ion transport” (GO:0006816), circadian rhythm and its regulation (GO:0007623, GO:0042752), “sensory system development” (GO:0048880), “phospholipid catabolic process” (GO:0009395), etc. (see Supplemental File S4 for the full list). The Reactome pathway “transcriptional regulation of granulopoiesis” (R-DME-9616222) was significantly overrepresented (adjusted *p* = 0.006). There were 82 PDD-associated *XtX* outlier SNPs mapped to 22 genes, including the clock genes *Clk*, *per*, and *wdb* (Figure 3D).

#### Voltinism

813 SNPs in 167 genes were found to be significantly associated with average voltinism (Table 1; Figure 3B; Figure 4). Top representative GO terms included “protein import” (GO:0017038), “imaginal disc-derived appendage development” (GO:0048737), “locomotor rhythm” (GO:0045475), etc. (see Supplemental File S4 for the full list). We wanted to see whether there might be any biological process under selection for voltinism that was not involved in diapause.

**Figure 4.**
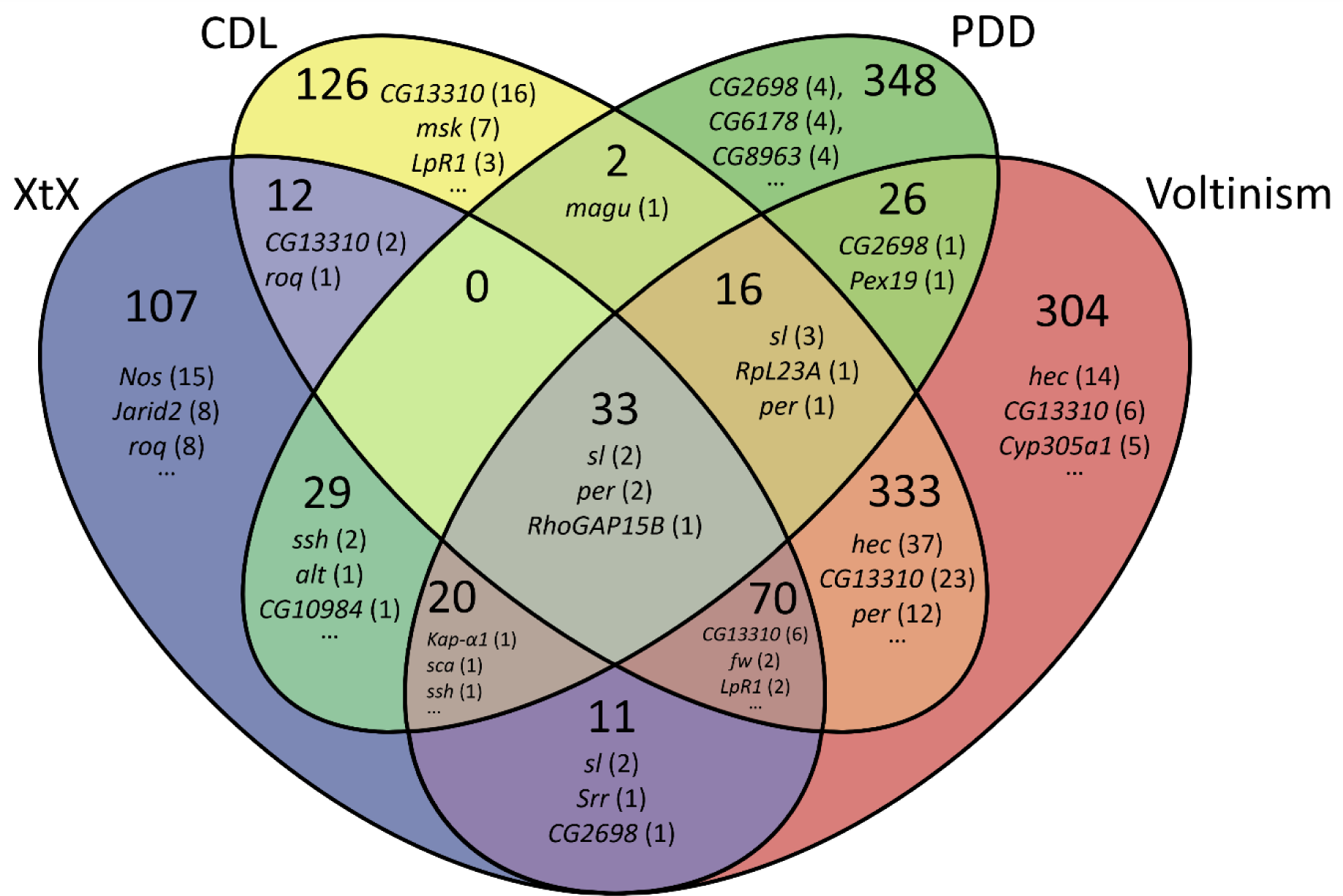
Venn diagram showing the number of unique and shared SNPs significant for *XtX* or association with voltinism, critical daylength (CDL), or post-diapause development (PDD) time. Up to 3 genes and their corresponding number of SNPs in parentheses are listed for each partition. See Supplemental File S3 for the full gene list.

At the SNP level, there were 4 *XtX* outliers (that mapped to a gene) uniquely associated with voltinism (Figure 4). However, at the gene level, any genes that were uniquely associated with voltinism (Supplemental File S3) did not contain any *XtX* outlier SNPs, and there were no GO terms enriched in these unique voltinism genes.

#### Pleiotropy

We identified 33 SNPs that were *XtX* outliers and significantly associated with all three phenology-related covariates (voltinism, CDL, and PDD). Only 5 of them were mapped to a *D. melanogaster* homolog: 1 in the gene *RhoGAP15B*, 2 in *sl*, and 2 in *per* (Figure 4, Figure 5).

**Figure 5.**
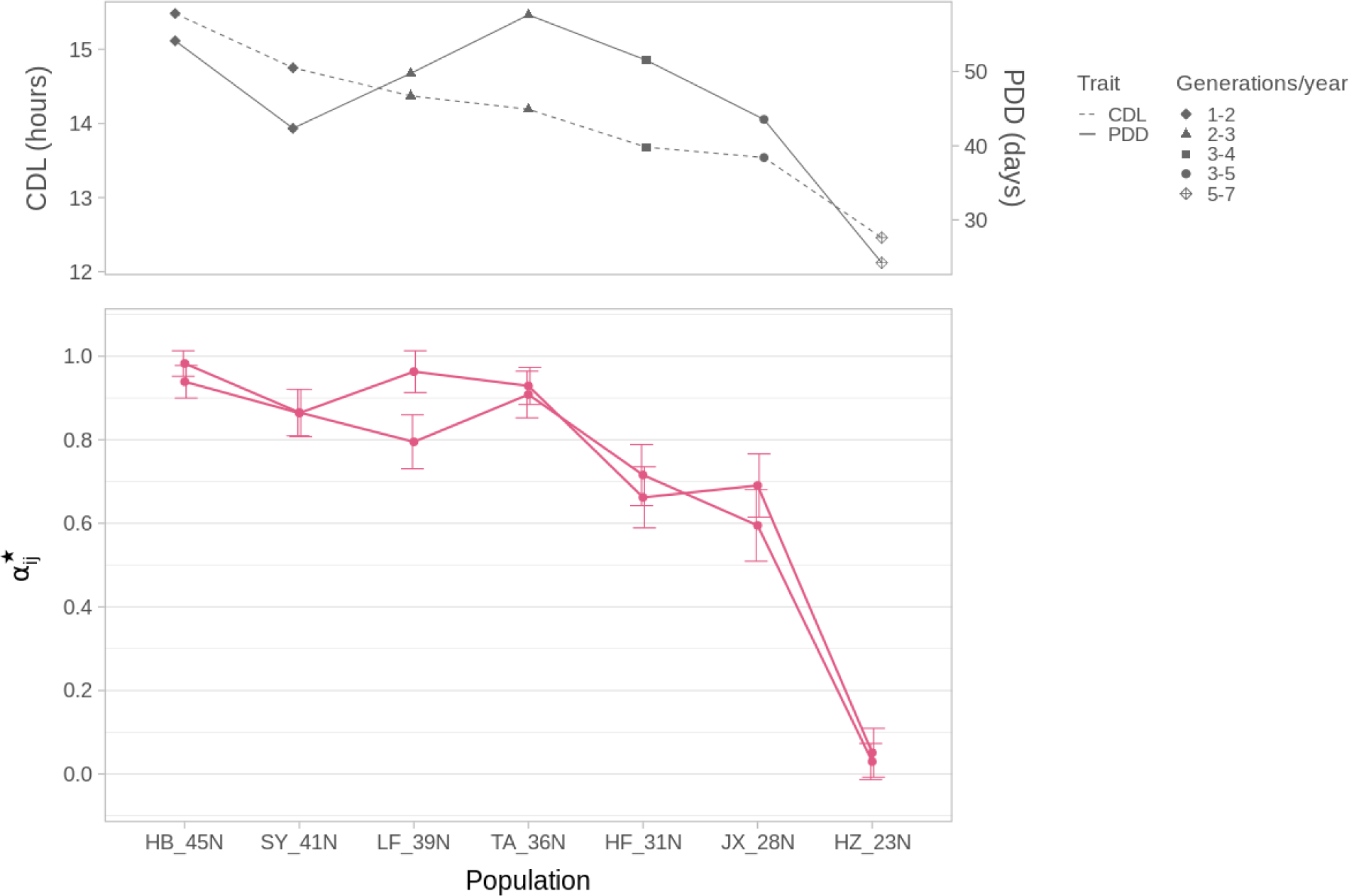
Population allele frequencies of *XtX* outliers in the core circadian clock gene *per*. Two *per* SNPs significantly associate with all three phenology covariates (voltinism, critical daylength (CDL), and post-diapause development (PDD) time). Top panel: covariates are plotted for comparison. Bottom panel: mean and standard deviation of the posterior distribution of the BayPass parameter β^*^_*ij*_, a proxy of allele frequency but with an untruncated Gaussian distribution (Gautier et al., 2010), plotted against populations arranged by latitude (left, northernmost; right, southernmost). Error bars (standard deviation) are jittered to avoid overlap. Overall trend is similar to that of allele frequencies calculated from raw read counts (data not shown).

Interestingly, out of the remaining 28 uncharacterized SNPs, 21 mapped to the same scaffold (NW_021136372.1, with 6 of the SNPs mapping to the predicted ACB genes “uncharacterized LOC114362050” and “uncharacterized LOC114362051” without *Drosophila* homologs). This is a relatively small scaffold (126 kb) putatively autosomal (Chromosome 12).

#### Other clinal genes under selection

Among latitudinally varying genes (associated with the latitude covariate; Supplemental File S3) that showed strong signals of selection (*XtX* outliers), *Jarid2* was the only clinal gene that was not associated with any of the three phenology-related covariates (Figure 3A; Supplemental File S3).

### Signatures of introgression between ACB and ECB

We used two four-taxon test statistics to detect signatures of introgression between ACB and ECB, particularly at the identified candidate loci above. We only used SNPs that were fixed in the outgroup (ALB) to polarize the sites with a final total of 5,551,517 usable biallelic SNPs. We calculated the *D*-statistic to estimate the extent of introgression genome-wide and calculated *f_d_* to estimate the proportion of introgression in small genomic windows.

For the tree topologies that were (((ECB_EA, ECB_PY), ACB), ALB), we found *D* > 0 for when any of the seven ACB populations was used as P_3_, though not all of the *D* values were significant (range of *D*: 1.67×10^-3^ to 2.58×10^-3^; range of *z*-score: 2.15 to 3.29; Figure 6A left panel). This suggests that there may have been more P_2_-P_3_ introgression (ECB_PY and ACB) compared to P_1_-P_3_ (ECB_EA and ACB). On the other hand, *D* values calculated for the tree topologies with ACB populations as ingroups showed that JX_28N had the most putative introgression with ECB relative to the other ACB populations (Figure 6A middle panel) while HZ_23N had the least (Figure 6A right panel).

**Figure 6.**
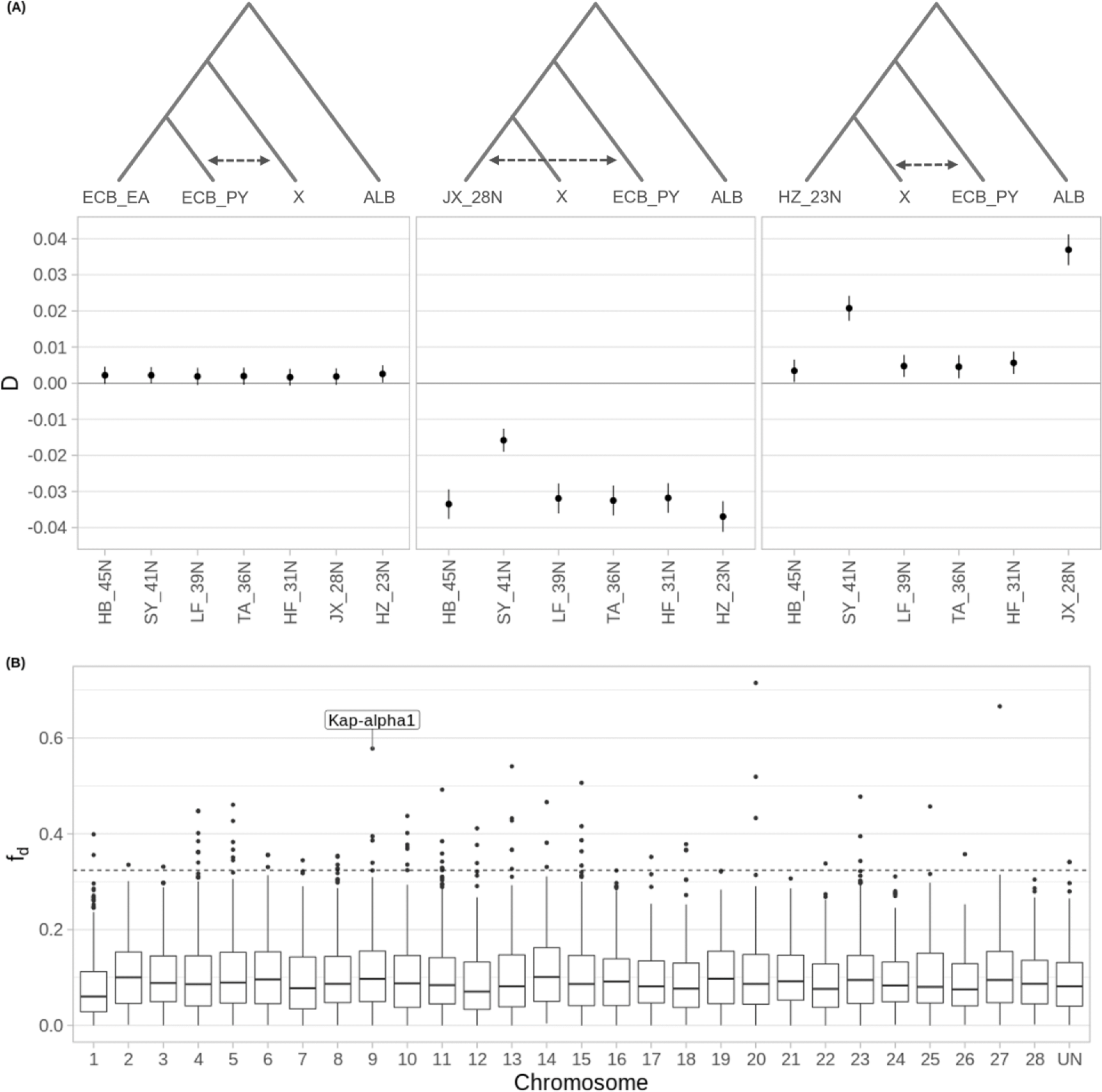
Four-taxon tests for signatures of introgression. **(A)** The 3 groups of tree topologies tested (top) and the corresponding whole-genome *D* (bottom) with 99.73% confidence interval (roughly equivalent to a *z*-score of 3). ACB populations used in the “X” position in the trees (top) are plotted along the x-axis (bottom). Dashed arrow (top) indicates the overall direction of introgression indicated by the *D* values (bottom). Left panel: ECB_PY showed more evidence of introgression than ECB_EA with the 7 ACB populations (all *D* > 0, not all |*z*| ≥ 3). Middle panel: JX_28N, the second southernmost population, showed relatively more evidence of introgression with ECB_PY than any of the other 6 ACB populations (all *D* < 0, all |*z*| > 3). Right panel: HZ_23N, the southernmost population, shows the least introgression with ECB_PY than any of the other 6 ACB populations (all *D* > 0, all |*z*| > 3). **(B)** *f_d_* calculated in genomic windows of 300 SNPs for the tree topology (((HZ_23N, SY_41N), ECB_PY), ALB). The window containing the putatively adaptive SNP in *Kap-α1* (see also Figure 3, 4) is labeled. ACB reference scaffolds are aligned to the 28 *Bombyx mori* chromosomes. Chr 1: the Z (sex) chromosome. UN: unplaced scaffolds. Horizontal dashed line: the top 1% *f_d_* threshold for this tree topology.

To pinpoint genomic regions with elevated introgression potentially related to the adaptive diapause response, we calculated *f_d_* for the topologies (((HZ_23N, ACB), ECB_PY), ALB), with HZ_23N in the P_1_ position as the “baseline” level of shared variation with ECB. In the 6 trees tested, with each of the remaining 6 ACB populations in the P_2_ position, we found that *Kap-α1* showed signatures of both introgression with ECB (*f_d_* outlier) and selection (*XtX* outlier). More specifically, *Kap-α1* was in an *f_d_* outlier window (top 1%) for the two trees with HB_45N and SY_41N in the P_2_ position (Figure 6; Supplemental Table S2).

**Table 2.**
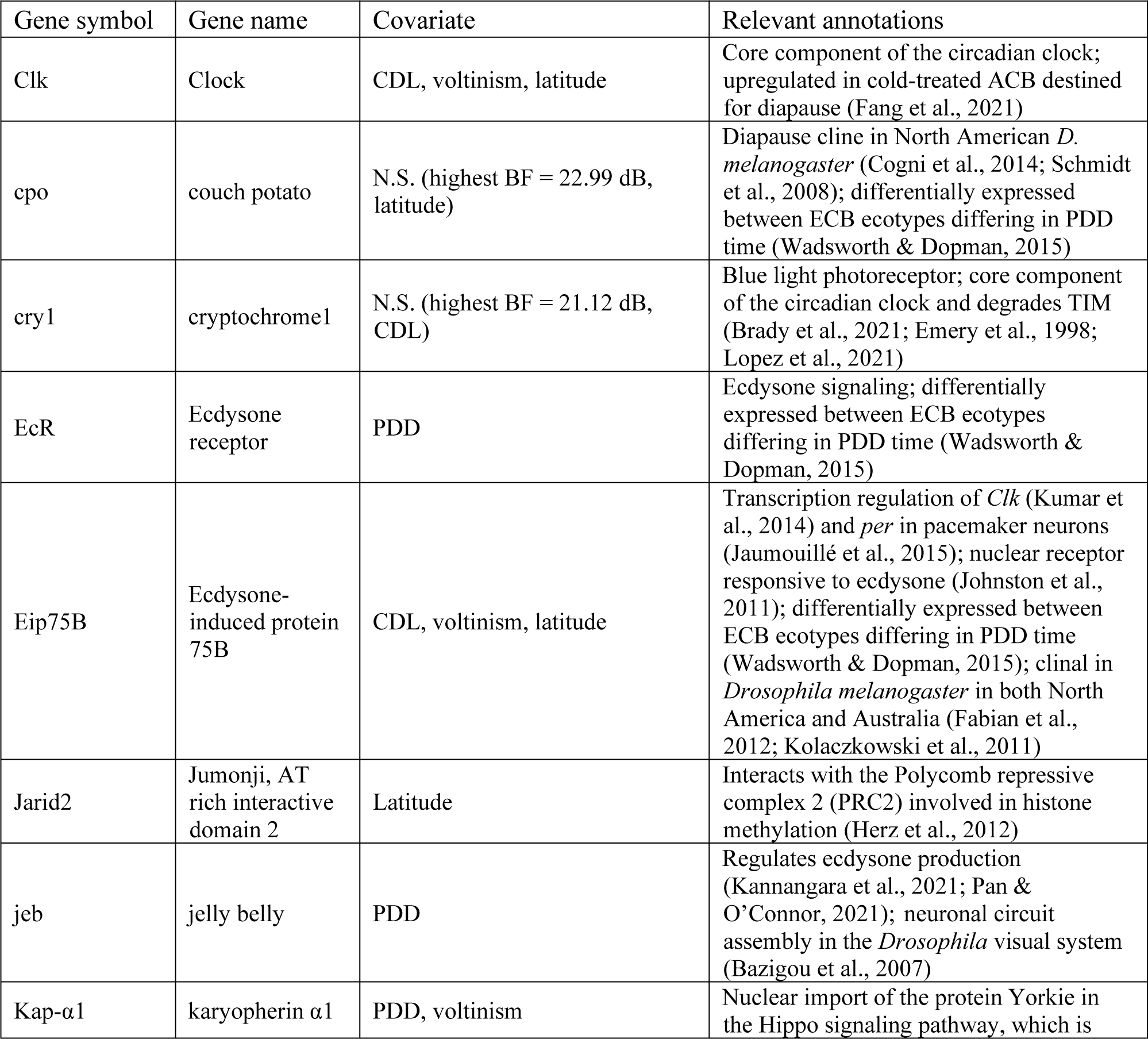
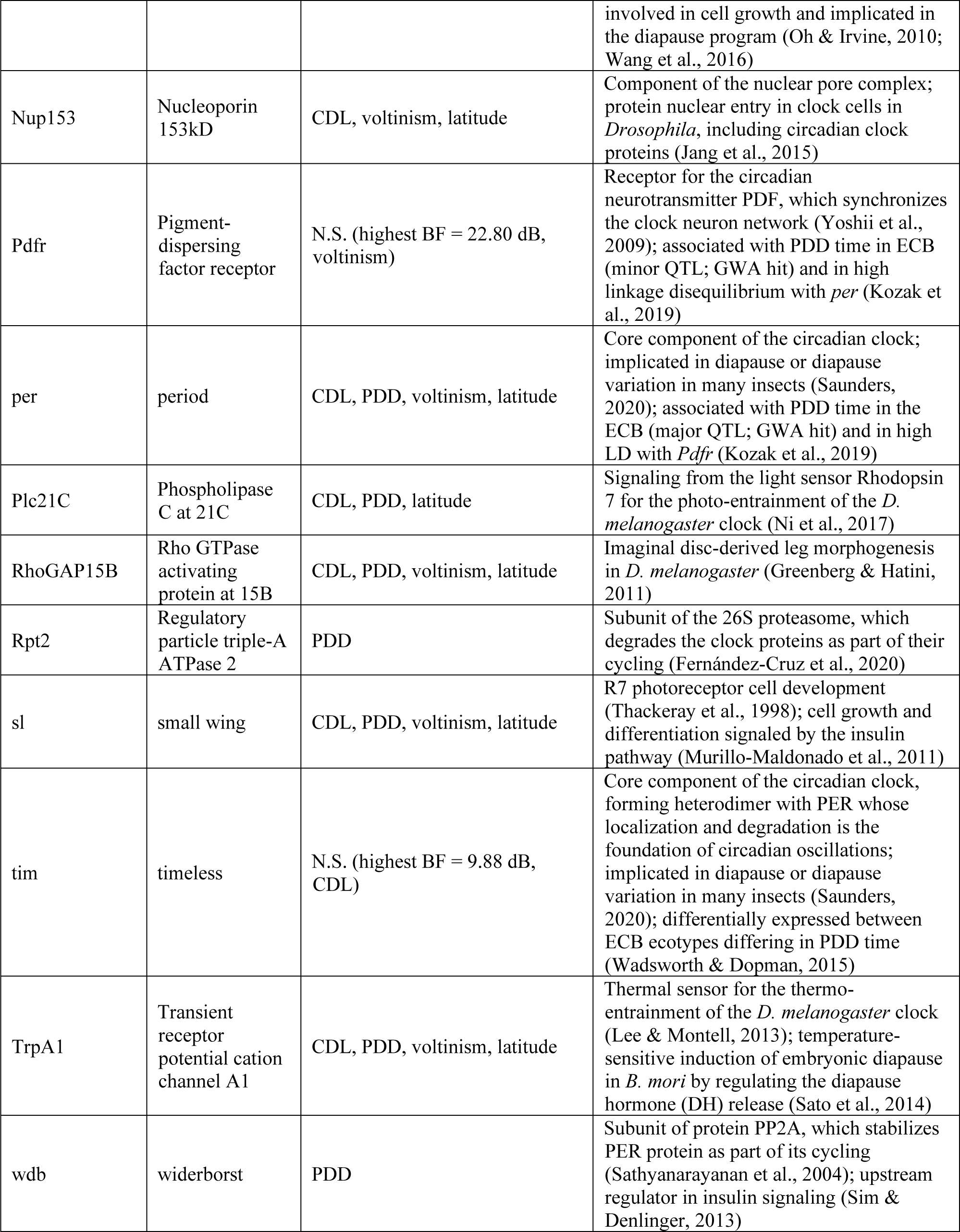
Genes mentioned in text. All genes follow Drosophila melanogaster nomenclature except cry1 (orthologous to cry in D. melanogaster). N.S.: not significant.

To make sure that the high *f_d_* resulted from introgression rather than ancestral structure (Eriksson & Manica, 2012; Martin et al., 2015; J. Smith & Kronforst, 2013), we calculated *d_xy_* and found that, consistent with introgression, *f_d_* outlier windows did not have higher *d_xy_* than the rest of the genome (Supplemental Figure S2). Still, *D* and *f_d_* tend to be inflated in regions of low *d_xy_* even without introgression (Martin et al., 2015), and other methods explicitly incorporating *d_xy_* to detect introgression have been proposed (Pfeifer & Kapan, 2019).

We found no evidence of recent introgression at *per*, the gene seemingly reused by ACB and ECB during phenological adaptation. To investigate ancestral standing variations as an alternative source of variation, we selected outlier SNPs identified in ACB (this study) and in ECB (Kozak et al., 2019) and checked for their alleles in the outgroup in our aligned data. We found that most of these SNPs were monomorphic in the outgroup, and the alternative allele would exclusively exist in only one of the two corn borer species where it was putatively under selection (Supplemental Table S3). That is to say, ACB and ECB did not share any derived alleles implicated in diapause at *per*.

As *f_d_* is more sensitive to introgression from P_3_ (ECB_PY) to P_2_ (ACB) than P_2_ to P_3_ (Martin et al., 2015), it is possible that adaptive introgression in the ACB to ECB direction could be better detected by using the two ECB as the ingroups and the ACB populations as P_3_. However, we found that for the structure (((ECB, ECB), ACB), ALB), genes previously shown to be significantly associated with diapause or voltine types in ECB, most notably *per*, *Pdfr*, and *cry1* (Kozak et al., 2019; Levy et al., 2015), did not have significantly elevated *f_d_* regardless of tree topology.

## DISCUSSION

Predicting evolutionary response to new environmental challenges is a central goal of modern evolutionary biology. Accumulating studies suggest that closely related species might be more likely to respond similarly at the genetic level to a novel environment because of shared access to ancestral variations and genetic background, and the more permeable reproductive barrier also makes it more likely for them to exchange adaptive alleles. Here, we used the parallel phenological evolution of the Asian and European corn borers to assess the repeatability of the underlying genetic evolution and the source of any reused genetic variations. We sampled 7 ACB populations spanning 23 degrees of latitude and found that among hundreds of candidate loci, *per* was repeatedly reused between ACB and ECB, but its alleles seem to have a more ancient source than recent introgression.

### Candidate genes and pathways in ACB

Perhaps the strongest candidate genes for phenology were *per*, *RhoGAP15B*, and *sl*. All three genes harbored SNPs potentially pleiotropic for voltinism, CDL, and PDD, with strong signatures of selection. Unlike the circadian clock gene *per* (discussed below), *RhoGAP15B* and *sl* have not been implicated in diapause or phenological adaptation in other species to our knowledge. In *D. melanogaster*, *RhoGAP15B* is involved in imaginal disc-derived leg morphogenesis (Greenberg & Hatini, 2011) and *sl* is necessary for proper eye development and growth (Murillo-Maldonado et al., 2011; Thackeray et al., 1998). Whether and how they affect diapause remains to be investigated. Generally speaking, a candidate gene could be underlying an unmeasured covarying trait rather than diapause. For example, the ECB ecotype that diapauses later in the year also has higher cold shock survival than the early-diapausing ecotype (Wadsworth et al., 2020). Such a trait would be expected to covary with shifts in diapause timing along the cline.

We also highlight here other interesting candidates for diapause with relevant known functions, albeit some with a weaker signature of selection. The first group of genes presented here is related to the circadian rhythm. In almost a century since Bünning’s foundational work in which he suggested the circadian clock as the basis of photoperiodism (Bünning, 1936) , there has been growing evidence supporting this hypothesis in plants (Sawa et al., 2008; Song et al., 2010) and insects (Denlinger et al., 2017, but see Emerson et al., 2009 and Saunders, 2010). Although the circadian component of photoperiodism seems to be strain- and temperature-dependent in ECB (Beck, 1989; Skopik & Bowen, 1976; Takeda & Skopik, 1985) and lacking in at least one population of ACB (i.e. JX_28N; Yang et al., 2014), recent genetic evidence suggests that the circadian clock does play an important role in adaptive circannual timing within the *Ostrinia* genus. In ECB, the clock genes *per* and *Pdfr* are major QTLs for diapause induction and/or termination, and PDD time is coupled with an intrinsic property of the circadian clock – the period of free-running activity rhythm (Kozak et al., 2019). In ACB, we found that the circadian clock pathway was overrepresented among candidate genes under latitudinally-varying selection. Therefore, even though the mechanistic model of clock-based photoperiodism is not yet well understood in corn borers (Saunders, 2012, 2016; Yang et al., 2014), genes that modulate or maintain functions of the circadian clock are especially interesting candidates for diapause in ACB. Specifically, we propose candidate genes related to the photo- and thermo-entrainment of the circadian clock (*Plc21C* and *TrpA1*, respectively), core components of the clock (*Clk*, in addition to *per*), nuclear import of the clock proteins (*Nup153*, *Kap-α1*), and other components of the clock machinery (*wdb*, *Rpt2*, *Eip75B*) (Table 2).

Among these genes, *Kap-α1* (also known as *importin α1*, or *impα1*) showed evidence of adaptive introgression. Its protein product IMPα1 is responsible for the nuclear translocation of TIM (and consequently PER) in *D. melanogaster* clock cells by binding to the NLS sequence of TIM (Jang et al., 2015). The delay between the transcription and nuclear translocation of PER and TIM is thought to be critical for the negative feedback loop and circadian oscillations (Curtin et al., 1995; Saez et al., 2011). However, IMPα1 expression did not seem to alter periodicity in *D. melanogaster* (Jang et al., 2015). Another possible explanation for *Kap-α1*’s correlation with PDD could be its involvement in the Hippo signaling pathway, which regulates cell growth and organ size and is one of numerous signaling pathways implicated in the diapause program (Y. Chen et al., 2017; Denlinger, 2022; N. Li et al., 2019; Oh & Irvine, 2010; Ragland et al., 2019; S. Wang et al., 2016).

The second group of interesting candidate genes is related to ecdysone signaling. Ecdysone signaling regulates developmental stage transitions and metamorphosis (Riddiford, 1993) and is central to transitions among stages of diapause (Denlinger, 2002). Ecdysone production in the prothoracic gland is thought to be stimulated by the prothoracicotropic hormone (PTTH) whose release in turn is regulated by the circadian clock (Denlinger, 2022). In many insects, the decrease of PTTH and ecdysone is linked to the induction of diapause, whereas their increase is linked to diapause termination (Denlinger, 2022; Denlinger et al., 2012; Gelman et al., 1992; Peypelut et al., 1990). We identify two genes involved in this process as candidates for diapause: *EcR* and *jeb*, both associated with PDD and not CDL. *EcR*, along with other nuclear receptors including *Eip75B*, modulates the coordinated activation and repression of transcription of many genes targeted by ecdysone (Johnston et al., 2011; K. Li et al., 2016). The gene *jeb* has an effect on, though not necessary for, ecdysone production through PTTH signaling (Kannangara et al., 2021; Pan & O’Connor, 2021). In ECB, *EcR* is upregulated in the strain that terminates diapause earlier compared to the late-terminating strain (Wadsworth & Dopman, 2015). It would be interesting to investigate whether *EcR* and *jeb* are upregulated in concert with the increase of ecdysone associated with diapause termination in ACB.

An important note on candidate genes is that some of the strongest candidates under selection in our data could not be functionally annotated by sequence homology. For example, “uncharacterized LOC114362050” on the scaffold NW_021136372.1 has numerous *XtX* outliers that are associated with both CDL and PDD (Supplemental File S3). While the ACB and ECB genomes are increasingly complete (Fang et al., 2021), future effort to characterize putatively important genic regions like LOC114362050 is much needed for a complete picture of the genetic basis of ecologically important traits.

### Genetic architectures

It is curious to note where the genetic architectures of CDL and PDD did and did not overlap at the gene level (Table 2). For example, *Eip75B* represses *EcR* in ecdysone signaling during developmental transition (Johnston et al., 2011); *Eip75B* has also been shown to directly interact with PER as well as *Clk* to regulate their expression (Jaumouillé et al., 2015; Kumar et al., 2014) , where *Clk*-mediated negative transcriptional feedback loop of *per* is the basis of circadian oscillations. Despite this network of interactions, only *per* was associated with both CDL and PDD in ACB. *Eip75B* and *Clk* covaried with CDL, whereas ecdysone signaling gene *EcR* (and *jeb*) covaried with PDD as mentioned above. There seems to be differential response to selection in interacting gene networks. In fact, CDL and PDD shared less than 10% of candidate SNPs (n = 51) (Figure 4) and roughly 25% of candidate genes (n = 21, genes with *D. melanogaster* homologs only) (Supplemental File S3). This is comparable with a previous study in *Rhagoletis pomonella*, in which the genetic architectures of initial diapause intensity and diapause termination were largely independent and overlapped on one of the chromosomes (Ragland et al., 2017). In our case, the shared candidate SNPs clustered in a handful of highly differentiated regions on a few chromosomes, especially on the Z chromosome where *per* and *sl* are located (Supplemental Figure S1). Overall, it seems that there is a low to intermediate level of genetic pleiotropy. A lower level of pleiotropy is considered to pose less constraint on the evolution of multiple traits required for adaptation; moreover, pleiotropy is increasingly recognized to even play a potentially conducive role in evolution (Chevin et al., 2010; Hansen, 2003; Hansen & Pélabon, 2021; Hughes & Leips, 2017; Ragland et al., 2017; Rennison & Peichel, 2022).

The importance of the diapause stage in phenology can be reflected in the fact that all highly differentiated genes associated with voltinism were also associated with one or both of the diapause traits (Figure 4). Conceivably, there could be other life history traits subject to selection during the growing season affecting variation in voltinism. For example, there could be adaptive thermal plasticity in the rate of development (Kong et al., 2019; Posledovich et al., 2014; Shama et al., 2011; Yamahira et al., 2007), whereby differential thermal optima for development affect generation time and potentially the number of generations. However, in both corn borer species, previous studies found that differences between populations in the thermal reaction norms of development rate and other life history traits were small and did not seem to correlate with voltine types (Calvin et al., 1991; Xiao et al., 2016). Furthermore, even when considering those genes that were only weakly differentiated between populations, no biological process stood out as being uniquely enriched for voltinism. Therefore, at least at the gene level, the genetic architecture of voltinism seems to be a combination of that of CDL and PDD.

### Repeated use of per

The circadian clock gene *per* was the only gene we found to be repeatedly used in the parallel evolution of ECB and ACB. Only at the gene level, that is. Indeed, it seems that a completely different set of SNPs underlies ACB’s cline. Perhaps even more surprisingly, most of these SNPs seemed to be new mutations (Supplemental Table S3).

The formation of the clines on the continents had been rapid. ACB and ECB made the host shift to maize independently around 500 years ago (Alexandre et al., 2013; Calcagno et al., 2017).

The ECB cline in North America formed within 100 years since its introduction around 1920 (Showers et al., 1975; Sparks & Young, 1971). The history of the ACB cline is less clear, but the adoption of maize cultivation over time across China was documented in historical records (S. Chen & Kung, 2016). By 1800 (within 250 years) all seven regions that we sampled from were cultivating maize (Figure 3 in S. Chen & Kung, 2016). Preexisting genetic variations (standing or introgressed) are thought to facilitate such rapid evolution (Barrett & Schluter, 2008) and especially parallel evolution (Bohutínská et al., 2021) . However, we did not find any evidence of recent introgression at *per*. The absence of most of the outlier SNPs in the outgroup ALB and of some in ECB’s sibling species *O. scapulalis* (preliminary, Supplemental Table S3) suggests independent mutational origins at *per* between the two corn borers.

In light of our findings at *per*, it is not surprising that *Pdfr* did not seem to contribute to adaptation in ACB, against our *a priori* expectation. In ECB, *per* is epistatic to the *Pdfr*- containing QTL for PDD, only affecting the phenotype when paired with one *per*-containing QTL allele (Kozak et al., 2019). The different set of SNPs at *per* in ACB likely altered the interaction with and selection dynamics at *Pdfr*. The case of *Pdfr* illustrates the importance of genetic interaction/background in the probability of gene reuse (Conte et al., 2012; Stern & Orgogozo, 2009).

On the other hand, *per* appears to be a frequent target of selection as seen in the corn borers and other systems (Cruciani et al., 2008; Kyriacou et al., 2008; Paolucci et al., 2016; Pruisscher et al., 2018). A possibility is that it is the circadian clock pathway that is targeted by selection when it comes to adaptation to seasonality. Specifically, another core clock gene, *tim*, is also found to underlie seasonal timing traits in several insects (Mathias et al., 2005; Pruisscher et al., 2018; Sandrelli et al., 2007). Reuse of pathways rather than genes is not uncommon across the tree of life. In plants, Arctic adaptation of three Brassicaceae species was found to involve more parallelism in stress response pathways rather than genes (Birkeland et al., 2020). *Arabidopsis*’s parallel adaptation to Cape Verde Islands involved earlier flowering time and was attributed to two genes used separately on two islands (Fulgione et al., 2022): *FRI* and *FLC*, two molecularly interacting genes that determine flowering time (Takada et al., 2019). In mammals, the *Mc1r* gene and its antagonist *agouti*, out of many pigmentation genes, seem to be preferred by selection for coat coloration (Gompel & Prud’homme, 2009) – in some cases both genes (Steiner et al., 2007) and in other cases *Mc1r* or *agouti* alone (Jones et al., 2018; Nachman et al., 2003).

Still, some genes seem to be reused more than other genes from the same pathway (Conte et al., 2012), which leads us back to the question of why *per* (and *tim*) is favored by selection.

Currently the ready explanation is that *per* has a large effect size on the timing traits – in ECB, the *per*-containing major QTL alone explains ∼30% of variance in PDD (Kozak et al., 2019; personal communication). The large effect size of *per* is corroborated by genetic crosses for diapause induction in the speckled wood butterfly *Pararge aegeria*, although in this system the *tim*-containing region seems to have a larger effect than *per* (Pruisscher et al., 2018). Other possible contributing factors (Conte et al., 2012) require further investigation, e.g. whether *per* has a higher mutation rate. Amino acid substitution rate at *per* was found to be relatively high (Regier et al., 1998). Length polymorphism of repetitive regions in *per* and other clock genes, presumably having a high mutation rate, has been noted as a substrate for evolution in *Drosophila* and birds (Cruciani et al., 2008; Kyriacou et al., 2008).

### Adaptive introgression at Kap-α1

Our results provide the first evidence for adaptive introgression between ACB and ECB. Specifically, we inferred adaptive introgression at *Kap-α1*. As we described above, how *Kap-α1* might contribute to the diapause phenotypes warrants further study. Importantly, this result is based on the global pattern of population differentiation. Therefore, there might be other introgressed adaptive genes missed by this analysis. One, there could be introgressed genes that are only locally adaptive, differing in allele frequency in just one population and not detected by our *XtX* outlier test. Similarly, there are introgressed genes not significantly differentiated among populations at our cutoff, but still more likely than not to be globally adaptive – for example, *Eip75B*. Three, the result could have been complicated by the choice of populations for the analysis. ECB likely went through bottlenecks when first introduced to North America and could be different from the Eurasian populations presumably more directly involved in introgression with ACB. Additionally, ancient introgression between ACB and ECB before the ACB populations split from each other is not detectable by the *f_d_* method.

We calculated the genome-wide average *D* which indicated that there was higher introgression from ECB to ACB than the opposite direction. This result contradicts previous studies using sympatric populations that found asymmetric gene flow from ACB to ECB and to ECB’s sibling species *O. scapulalis* (Bourguet et al., 2014; Y. Wang et al., 2017). Understanding of adaptive introgression between ACB and ECB would benefit from future studies incorporating Eurasian ECB populations and haplotype information.

## Conclusion

Repeated use of large effect loci like *per* notwithstanding, the genetic basis of parallel evolution proves to be complicated. This is partly due to the complex genetic architecture of traits like diapause and partly due to the evolutionary history unique to lineages (genetic background, demographic history, etc.). Nonetheless, there appears to be coordinated evolution of traits and of interacting genes and pathways. Understanding the intricate details in the interaction of environment, genes and biological context in individual cases is the critical step before we can generalize to a unifying theory for the prediction of evolution.

## Acknowledgments

We thank the funding agency National Science Foundation (DEB-1257251 to E.B.D.) and Tufts University (to E.B.D.). We thank Dr. Genevieve Kozak (University of Massachusetts Dartmouth) for discussions and sharing of analysis pipeline.

## REFERENCES

Adrion, J. R., Hahn, M. W., & Cooper, B. S. (2015). Revisiting classic clines in Drosophila melanogaster in the age of genomics. Trends in Genetics : TIG, 31(8), 434–444. https://doi.org/10.1016/j.tig.2015.05.006

Alexandre, H., Ponsard, S., Bourguet, D., Vitalis, R., Audiot, P., Cros-Arteil, S., & Streiff, R. (2013). When History Repeats Itself: Exploring the Genetic Architecture of Host-Plant Adaptation in Two Closely Related Lepidopteran Species. PLoS ONE, 8(7). https://doi.org/10.1371/journal.pone.0069211

Altschul, S. F., Gish, W., Miller, W., Myers, E. W., & Lipman, D. J. (1990). Basic local alignment search tool. Journal of Molecular Biology, 215(3), 403–410. https://doi.org/10.1016/S0022-2836(05)80360-2

Ammar, R. (2019). *RandomcoloR: Generate Attractive Random Colors* (R package version 1.1.0.1) [Computer software]. https://CRAN.R-project.org/package=randomcoloR

Ammar, R. (2020). *forcats: Tools for Working with Categorical Variables (Factors)* (R package version 0.5.0) [Computer software]. https://CRAN.R-project.org/package=forcats

Andrews, S. (2010). FastQC: A Quality Control Tool for High Throughput Sequence Data [Online]. http://www.bioinformatics.babraham.ac.uk/projects/fastqc/

Anduaga, A. M., Nagy, D., Costa, R., & Kyriacou, C. P. (2018). Diapause in Drosophila melanogaster— Photoperiodicity, cold tolerance and metabolites. Journal of Insect Physiology, 105, 46–53. https://doi.org/10.1016/j.jinsphys.2018.01.003

Arendt, J., & Reznick, D. (2008). Convergence and parallelism reconsidered: What have we learned about the genetics of adaptation? Trends in Ecology & Evolution, 23(1), 26–32. https://doi.org/10.1016/j.tree.2007.09.011

Barrett, R. D. H., & Schluter, D. (2008). Adaptation from standing genetic variation. Trends in Ecology & Evolution, 23(1), 38–44. https://doi.org/10.1016/j.tree.2007.09.008

Batz, Z. A., Clemento, A. J., Fritzenwanker, J., Ring, T. J., Garza, J. C., & Armbruster, P. A. (2020). Rapid adaptive evolution of the diapause program during range expansion of an invasive mosquito. Evolution; International Journal of Organic Evolution, 74(7), 1451–1465. https://doi.org/10.1111/evo.14029

Bazigou, E., Apitz, H., Johansson, J., Lorén, C. E., Hirst, E. M. A ., Chen, P.-L., Palmer, R. H., & Salecker, I. (2007). Anterograde Jelly belly and Alk Receptor Tyrosine Kinase Signaling Mediates Retinal Axon Targeting in Drosophila. Cell, 128(5), 961–975. https://doi.org/10.1016/j.cell.2007.02.024

Bean, D. W., Dalin, P., & Dudley, T. L. (2012). Evolution of critical day length for diapause induction enables range expansion of Diorhabda carinulata, a biological control agent against tamarisk (Tamarix spp.). Evolutionary Applications, 5(5), 511–523. https://doi.org/10.1111/j.1752-4571.2012.00262.x

Beck, S. D. (1983). Thermal and thermoperiodic effects on larval development and diapause in the European corn borer, Ostrinia nubilalis. Journal of Insect Physiology, 29(1), 107–112. https://doi.org/10.1016/0022-1910(83)90113-0

Beck, S. D. (1989). Factors influencing the intensity of larval diapause in Ostrinia nubilalis. Journal of Insect Physiology, 35(2), 75–79. https://doi.org/10.1016/0022-1910(89)90039-5

Birkeland, S., Gustafsson, A. L. S., Brysting, A. K., Brochmann, C., & Nowak, M. D. (2020). Multiple Genetic Trajectories to Extreme Abiotic Stress Adaptation in Arctic Brassicaceae. Molecular Biology and Evolution, 37(7), 2052–2068. https://doi.org/10.1093/molbev/msaa068

Bohutínská, M., Vlček, J., Yair, S., Laenen, B., Konečná, V., Fracassetti, M., Slotte, T., & Kolář, F. (2021). Genomic basis of parallel adaptation varies with divergence in Arabidopsis and its relatives. Proceedings of the National Academy of Sciences, 118(21), e2022713118. https://doi.org/10.1073/pnas.2022713118

Boitard, S., Schlötterer, C., Nolte, V., Pandey, R. V., & Futschik, A. (2012). Detecting Selective Sweeps from Pooled Next-Generation Sequencing Samples. Molecular Biology and Evolution, 29(9), 2177–2186. https://doi.org/10.1093/molbev/mss090

Bolger, A. M., Lohse, M., & Usadel, B. (2014). Trimmomatic: A flexible trimmer for Illumina sequence data. Bioinformatics, 30(15), 2114–2120. https://doi.org/10.1093/bioinformatics/btu170

Bourguet, D., Ponsard, S., Streiff, R., Meusnier, S., Audiot, P., Li, J., & Wang, Z.-Y. (2014). ‘Becoming a species by becoming a pest’ or how two maize pests of the genus Ostrinia possibly evolved through parallel ecological speciation events. Molecular Ecology, 23(2), 325–342. https://doi.org/10.1111/mec.12608

Boyle, J. H., Rastas, P. M. A., Huang, X., Garner, A. G., Vythilingam, I., & Armbruster, P. A. (2021). A Linkage-Based Genome Assembly for the Mosquito Aedes albopictus and Identification of Chromosomal Regions Affecting Diapause. Insects, 12(2), 167. https://doi.org/10.3390/insects12020167

Bradshaw, W. E., & Holzapfel, C. M. (2010). Light, time, and the physiology of biotic response to rapid climate change in animals. Annual Review of Physiology, 72, 147–166. https://doi.org/10.1146/annurev-physiol-021909-135837

Brady, D., Saviane, A., Cappellozza, S., & Sandrelli, F. (2021). The Circadian Clock in Lepidoptera. Frontiers in Physiology, 12, 776826. https://doi.org/10.3389/fphys.2021.776826

Brownrigg, R., Minka, T. P., & Deckmyn, A. (2018). *maps: Draw Geographical Maps* (R package version 3.3.0) [Computer software]. https://CRAN.R-project.org/package=maps

Bünning, E. (1936). Die endogene Tagesrhythmik als Grundlage der photoperiodischen Reaktion. Berichte Der Deutschen Botanischen Gesellschaft, 54, 590–607.

Calcagno, V., Mitoyen, C., Audiot, P., Ponsard, S., Gao, G., Lu, Z., Wang, Z., He, K., & Bourguet, D. (2017). Parallel evolution of behaviour during independent host-shifts following maize introduction into Asia and Europe. Evolutionary Applications, 10(9), 881–889. https://doi.org/10.1111/eva.12481

Calvin, D. D., Higgins, R. A., Knapp, M. C., Poston, F. L., Welch, S. M., Showers, W. B., Witkowski, J. F., Mason, C. E., Chiang, H. C., & Keaster, A. J. (1991). Similarities in Developmental Rates of Geographically Separate European Corn Borer (Lepidoptera: Pyralidae) Populations. Environmental Entomology, 20(2), 441–449. https://doi.org/10.1093/ee/20.2.441

Cavedon, M., Gubili, C., Heppenheimer, E., vonHoldt, B., Mariani, S., Hebblewhite, M., Hegel, T., Hervieux, D., Serrouya, R., Steenweg, R., Weckworth, B. V., & Musiani, M. (2019). Genomics, environment and balancing selection in behaviourally bimodal populations: The caribou case. Molecular Ecology, 28(8), 1946–1963. https://doi.org/10.1111/mec.15039

Chen, H., & Boutros, P. C. (2011). VennDiagram: A package for the generation of highly-customizable Venn and Euler diagrams in R. BMC Bioinformatics, 12(1), 35. https://doi.org/10.1186/1471-2105-12-35

Chen, S., & Kung, J. K. (2016). Of maize and men: The effect of a New World crop on population and economic growth in China. Journal of Economic Growth, 21(1), 71–99. https://doi.org/10.1007/s10887-016-9125-8

Chen, Y., Jiang, T., Zhu, J., Xie, Y., Tan, Z., Chen, Y., Tang, S., Hao, B., Wang, S., Huang, J., & Shen, X. (2017). Transcriptome sequencing reveals potential mechanisms of diapause preparation in bivoltine silkworm Bombyx mori (Lepidoptera: Bombycidae). Comparative Biochemistry and Physiology Part D: Genomics and Proteomics, 24, 68–78. https://doi.org/10.1016/j.cbd.2017.07.003

Chen, Y.-S., Chen, C., He, H.-M., Xia, Q.-W., & Xue, F.-S. (2013). Geographic variation in diapause induction and termination of the cotton bollworm, Helicoverpa armigera Hübner (Lepidoptera: Noctuidae). Journal of Insect Physiology, 59(9), 855–862. https://doi.org/10.1016/j.jinsphys.2013.06.002

Chessel, D., Dufour, A. B., & Thioulouse, J. (2004). The ade4 package—I : One-table methods. R News, 4(1), 5–10.

Chevin, L.-M., Martin, G., & Lenormand, T. (2010). Fisher’s Model and the Genomics of Adaptation: Restricted Pleiotropy, Heterogenous Mutation, and Parallel Evolution. Evolution, 64(11), 3213–3231. https://doi.org/10.1111/j.1558-5646.2010.01058.x

Cogni, R., Kuczynski, C., Koury, S., Lavington, E., Behrman, E. L., O’Brien, K. R., Schmidt, P. S., & Eanes, W. F. (2014). The Intensity of Selection Acting on the Couch Potato Gene—Spatial–Temporal Variation in a Diapause Cline. Evolution, 68(2), 538–548. https://doi.org/10.1111/evo.12291

Colosimo, P. F., Hosemann, K. E., Balabhadra, S., Villarreal, G., Dickson, M., Grimwood, J., Schmutz, J., Myers, R. M., Schluter, D., & Kingsley, D. M. (2005). Widespread Parallel Evolution in Sticklebacks by Repeated Fixation of Ectodysplasin Alleles. Science, 307(5717), 1928–1933. https://doi.org/10.1126/science.1107239

Conte, G. L., Arnegard, M. E., Peichel, C. L., & Schluter, D. (2012). The probability of genetic parallelism and convergence in natural populations. Proc. R. Soc. B, 279(1749), 5039–5047. https://doi.org/10.1098/rspb.2012.2146

Cruciani, F., Trombetta, B., Labuda, D., Modiano, D., Torroni, A., Costa, R., & Scozzari, R. (2008). Genetic diversity patterns at the human clock gene period 2 are suggestive of population-specific positive selection. European Journal of Human Genetics, 16(12), 1526–1534. https://doi.org/10.1038/ejhg.2008.105

Curtin, K. D., Huang, Z. J., & Rosbash, M. (1995). Temporally regulated nuclear entry of the Drosophila period protein contributes to the circadian clock. Neuron, 14(2), 365–372. https://doi.org/10.1016/0896-6273(95)90292-9

Danilevskiĭ, A. S. (1965). *Photoperiodism and Seasonal Development of Insects*. Oliver & Boyd. Denlinger, D. L. (2002). Regulation of diapause. *Annual Review of Entomology*, *47*, 93–122. https://doi.org/10.1146/annurev.ento.47.091201.145137

Denlinger, D. L. (Ed.). (2022). Molecular Signaling Pathways that Regulate Diapause. In Insect Diapause (pp. 240–292). Cambridge University Press. https://doi.org/10.1017/9781108609364.010

Denlinger, D. L., Hahn, D. A., Merlin, C., Holzapfel, C. M., & Bradshaw, W. E. (2017). Keeping time without a spine: What can the insect clock teach us about seasonal adaptation? Philosophical Transactions of the Royal Society B: Biological Sciences, 372(1734), 20160257. https://doi.org/10.1098/rstb.2016.0257

Denlinger, D. L., Yocum, G. D., & Rinehart, J. P. (2012). 10—Hormonal Control of Diapause. In L. I. Gilbert (Ed.), Insect Endocrinology (pp. 430–463). Academic Press. https://doi.org/10.1016/B978-0-12-384749-2.10010-X

Dowle, E. J., Powell, T. H. Q., Doellman, M. M., Meyers, P. J., Calvert, M. B., Walden, K. K. O., Robertson, H. M., Berlocher, S. H., Feder, J. L., Hahn, D. A., & Ragland, G. J. (2020). Genome-wide variation and transcriptional changes in diverse developmental processes underlie the rapid evolution of seasonal adaptation. Proceedings of the National Academy of Sciences, 117(38), 23960–23969. https://doi.org/10.1073/pnas.2002357117

Dowle, M., & Srinivasan, A. (2020). *data.table: Extension of ‘data.fram’* (R package version 1.13.0) [Computer software]. https://CRAN.R-project.org/package=data.table

Durand, E. Y., Patterson, N., Reich, D., & Slatkin, M. (2011). Testing for Ancient Admixture between Closely Related Populations. Molecular Biology and Evolution, 28(8), 2239–2252. https://doi.org/10.1093/molbev/msr048

Emerson, K. J., Dake, S. J., Bradshaw, W. E., & Holzapfel, C. M. (2009). Evolution of photoperiodic time measurement is independent of the circadian clock in the pitcher-plant mosquito, Wyeomyia smithii. Journal of Comparative Physiology. A, Neuroethology, Sensory, Neural, and Behavioral Physiology, 195(4), 385–391. https://doi.org/10.1007/s00359-009-0416-9

Emery, P., So, W. V., Kaneko, M., Hall, J. C., & Rosbash, M. (1998). CRY, a Drosophila Clock and Light-Regulated Cryptochrome, Is a Major Contributor to Circadian Rhythm Resetting and Photosensitivity. Cell, 95(5), 669–679. https://doi.org/10.1016/S0092-8674(00)81637-2

Erickson, P. A., Weller, C. A., Song, D. Y., Bangerter, A. S., Schmidt, P., & Bergland, A. O. (2020). Unique genetic signatures of local adaptation over space and time for diapause, an ecologically relevant complex trait, in Drosophila melanogaster. PLOS Genetics, 16(11), e1009110. https://doi.org/10.1371/journal.pgen.1009110

Eriksson, A., & Manica, A. (2012). Effect of ancient population structure on the degree of polymorphism shared between modern human populations and ancient hominins. Proceedings of the National Academy of Sciences, 109(35), 13956–13960. https://doi.org/10.1073/pnas.1200567109

Fabian, D. K., Kapun, M., Nolte, V., Kofler, R., Schmidt, P. S., Schlötterer, C., & Flatt, T. (2012). Genome -wide patterns of latitudinal differentiation among populations of Drosophila melanogaster from North America. Molecular Ecology, 21(19), 4748–4769. https://doi.org/10.1111/j.1365-294X.2012.05731.x

Fang, G., Zhang, Q., Chen, X., Cao, Y., Wang, Y., Qi, M., Wu, N., Qian, L., Zhu, C., Huang, Y., & Zhan, S. (2021). The draft genome of the Asian corn borer yields insights into ecological adaptation of a devastating maize pest. Insect Biochemistry and Molecular Biology, 103638. https://doi.org/10.1016/j.ibmb.2021.103638

Feder, J. L., Berlocher, S. H., Roethele, J. B., Dambroski, H., Smith, J. J., Perry, W. L., Gavrilovic, V., Filchak, K. E., Rull, J., & Aluja, M. (2003). Allopatric genetic origins for sympatric host-plant shifts and race formation in Rhagoletis. Proceedings of the National Academy of Sciences, 100(18), 10314–10319. https://doi.org/10.1073/pnas.1730757100

Fernández -Cruz, I., Sánch ez-Díaz, I., Narváez -Padilla, V., & Reynaud, E. (2020). Rpt2 proteasome subunit reduction causes Parkinson’s disease like symptoms in Drosophila. IBRO Reports, 9, 65–77. https://doi.org/10.1016/j.ibror.2020.07.001

Frachon, L., Bartoli, C., Carrère, S., Bouchez, O., Chaubet, A., Gautier, M., Roby, D., & Roux, F. (2018). A Genomic Map of Climate Adaptation in Arabidopsis thaliana at a Micro-Geographic Scale. Frontiers in Plant Science, 9. https://doi.org/10.3389/fpls.2018.00967

French, B. W., Coates, B. S., & Sappington, T. W. (2014). Inheritance of an extended diapause trait in the Northern corn rootworm, Diabrotica barberi (Coleoptera: Chrysomelidae). Journal of Applied Entomology, 138(3), 213–221. https://doi.org/10.1111/j.1439-0418.2012.01751.x

Frolov, A. N., Bourguet, D., & Ponsard, S. (2007). Reconsidering the taxomony of several Ostrinia species in the light of reproductive isolation: A tale for Ernst Mayr. Biological Journal of the Linnean Society, 91(1), 49–72. https://doi.org/10.1111/j.1095-8312.2007.00779.x

Fulgione, A., Neto, C., Elfarargi, A. F., Tergemina, E., Ansari, S., Göktay, M., Dinis, H., Döring, N., Flood, P. J., Rodriguez-Pacheco, S., Walden, N., Koch, M. A., Roux, F., Hermisson, J., & Hancock, A. M. (2022). Parallel reduction in flowering time from de novo mutations enable evolutionary rescue in colonizing lineages. Nature Communications, 13(1), 1461. https://doi.org/10.1038/s41467-022-28800-z

Garcia-Moreno, A., López -Domínguez, R., Villatoro -García, J. A., Ramirez -Mena, A., Aparicio-Puerta, E., Hackenberg, M., Pascual-Montano, A., & Carmona-Saez, P. (2022). Functional Enrichment Analysis of Regulatory Elements. Biomedicines, 10(3), 590. https://doi.org/10.3390/biomedicines10030590

Gautier, M. (2015). Genome-Wide Scan for Adaptive Divergence and Association with Population-Specific Covariates. Genetics, 201(4), 1555–1579. https://doi.org/10.1534/genetics.115.181453

Gautier, M. (2019, November 22). *BayPass version 2.2 User Manual*. http://www1.montpellier.inra.fr/CBGP/software/baypass/files/BayPass_manual_2.2.pdf

Gautier, M., Foucaud, J., Gharbi, K., Cézard, T., Galan, M., Loiseau, A., Thomson, M., Pudlo, P., Kerdelhué, C., & Estoup, A. (2013). Estimation of population allele frequencies from next-generation sequencing data: Pool- versus individual-based genotyping. Molecular Ecology, 22(14), 3766–3779. https://doi.org/10.1111/mec.12360

Gautier, M., Hocking, T. D., & Foulley, J.-L. (2010). A Bayesian Outlier Criterion to Detect SNPs under Selection in Large Data Sets. PLOS ONE, 5(8), e11913. https://doi.org/10.1371/journal.pone.0011913

Gelman, D. B., Thyagaraja, B. S., Kelly, T. J., Masler, E. P., Bell, R. A., & Borkovec, A. B. (1992). Prothoracicotropic hormone levels in brains of the European corn borer, Ostrinia nubilalis: Diapause vs the non-diapause state. Journal of Insect Physiology, 38(5), 383–395. https://doi.org/10.1016/0022-1910(92)90063-J

Golczer, G. (2019). *Genetic basis and consequences of variation in seasonal timing.* [Doctoral dissertation, Tufts University]. ProQuest Dissertations and Theses Global.

Gompel, N., & Prud’homme, B. (2009). The causes of repeated genetic evolution. Developmental Biology, 332(1), 36–47. https://doi.org/10.1016/j.ydbio.2009.04.040

Green, R. E., Krause, J., Briggs, A. W., Maricic, T., Stenzel, U., Kircher, M., Patterson, N., Li, H., Zhai, W., Fritz, M. H.-Y., Hansen, N. F., Durand, E. Y., Malaspinas, A.-S., Jensen, J. D., Marques-Bonet, T., Alkan, C., Prüfer, K., Meyer, M., Burbano, H. A., … Pääbo, S. (2010). A Draft Sequence of the Neandertal Genome. Science, 328(5979), 710–722. https://doi.org/10.1126/science.1188021

Greenberg, L., & Hatini, V. (2011). Systematic expression and loss-of-function analysis defines spatially restricted requirements for Drosophila RhoGEFs and RhoGAPs in leg morphogenesis. Mechanisms of Development, 128(1), 5–17. https://doi.org/10.1016/j.mod.2010.09.001

Günther, T., & Coop, G. (2013). Robust Identification of Local Adaptation from Allele Frequencies. Genetics, 195(1), 205–220. https://doi.org/10.1534/genetics.113.152462

Hansen, T. F. (2003). Is modularity necessary for evolvability?: Remarks on the relationship between pleiotropy and evolvability. Biosystems, 69(2), 83–94. https://doi.org/10.1016/S0303-2647(02)00132-6

Hansen, T. F., & Pélabon, C. (2021). Evolvability: A Quantitative -Genetics Perspective. Annual Review of Ecology, Evolution, and Systematics, 52(1), 153–175. https://doi.org/10.1146/annurev-ecolsys-011121-021241

Herz, H.-M., Mohan, M., Garrett, A. S., Miller, C., Casto, D., Zhang, Y., Seidel, C., Haug, J. S., Florens, L., Washburn, M. P., Yamaguchi, M., Shiekhattar, R., & Shilatifard, A. (2012). Polycomb Repressive Complex 2-Dependent and -Independent Functions of Jarid2 in Transcriptional Regulation in Drosophila. Molecular and Cellular Biology, 32(9), 1683–1693. https://doi.org/10.1128/MCB.06503-11

Hivert, V., Leblois, R., Petit, E. J., Gautier, M., & Vitalis, R. (2018). Measuring Genetic Differentiation from Pool-seq Data. Genetics, 210(1), 315–330. https://doi.org/10.1534/genetics.118.300900

Hoffmann, A. A., & Sgrò, C. M. (2011). Climate change and evolutionary adaptation. Nature, 470(7335), 479–485. https://doi.org/10.1038/nature09670

Huang, L., Tang, J., Chen, C., He, H., Gao, Y., & Xue, F. (2020). Diapause incidence and critical day length of Asian corn borer (Ostrinia furnacalis) populations exhibit a latitudinal cline in both pure and hybrid strains. Journal of Pest Science, 93(2), 559–568. https://doi.org/10.1007/s10340-019-01179-5

Hughes, K. A., & Leips, J. (2017). Pleiotropy, constraint, and modularity in the evolution of life histories: Insights from genomic analyses. Annals of the New York Academy of Sciences, 1389(1), 76–91. https://doi.org/10.1111/nyas.13256

Jang, A. R., Moravcevic, K., Saez, L., Young, M. W., & Sehgal, A. (2015). Drosophila TIM Binds Importin α1, and Acts as an Adapter to Transport PER to the Nucleus. PLOS Genetics, 11(2), e1004974. https://doi.org/10.1371/journal.pgen.1004974

Jaumouillé, E., Machado Almeida, P., Stähli, P., Koch, R., & Nagoshi, E. (2015). Transcriptional Regulation via Nuclear Receptor Crosstalk Required for the Drosophila Circadian Clock. Current Biology, 25(11), 1502– 1508. https://doi.org/10.1016/j.cub.2015.04.017

Jeffreys, H. (1998). The Theory of Probability. OUP Oxford.

Johnston, D. M., Sedkov, Y., Petruk, S., Riley, K. M., Fujioka, M., Jaynes, J. B., & Mazo, A. (2011). Ecdysone- and NO-Mediated Gene Regulation by Competing EcR/Usp and E75A Nuclear Receptors during Drosophila Development. Molecular Cell, 44(1), 51–61. https://doi.org/10.1016/j.molcel.2011.07.033

Jones, M. R., Mills, L. S., Alves, P. C., Callahan, C. M., Alves, J. M., Lafferty, D. J. R., Jiggins, F. M., Jensen, J. D., Melo-Ferreira, J., & Good, J. M. (2018). Adaptive introgression underlies polymorphic seasonal camouflage in snowshoe hares. Science, 360(6395), 1355–1358. https://doi.org/10.1126/science.aar5273

Kannangara, J. R., Mirth, C. K., & Warr, C. G. (2021). Regulation of ecdysone production in Drosophila by neuropeptides and peptide hormones. Open Biology, 11(2), 200373. https://doi.org/10.1098/rsob.200373

Kassambara, A. (2020). *ggpubr: “ggplot2” Based Publication Ready Plots* (R package version 0.4.0) [Computer software]. https://CRAN.R-project.org/package=ggpubr

Kawamoto, M., Jouraku, A., Toyoda, A., Yokoi, K., Minakuchi, Y., Katsuma, S., Fujiyama, A., Kiuchi, T., Yamamoto, K., & Shimada, T. (2019). High-quality genome assembly of the silkworm, Bombyx mori. Insect Biochemistry and Molecular Biology, 107, 53–62. https://doi.org/10.1016/j.ibmb.2019.02.002

Kofler, R., Pandey, R. V., & Schlötterer, C. (2011). PoPoolation2: Identifying differentiation between populations using sequencing of pooled DNA samples (Pool-Seq). Bioinformatics, 27(24), 3435–3436. https://doi.org/10.1093/bioinformatics/btr589

Kolaczkowski, B., Kern, A. D., Holloway, A. K., & Begun, D. J. (2011). Genomic Differentiation Between Temperate and Tropical Australian Populations of Drosophila melanogaster. Genetics, 187(1), 245–260. https://doi.org/10.1534/genetics.110.123059

Kong, J. D., Hoffmann, A. A., & Kearney, M. R. (2019). Linking thermal adaptation and life-history theory explains latitudinal patterns of voltinism. Philosophical Transactions of the Royal Society B: Biological Sciences, 374(1778), 20180547. https://doi.org/10.1098/rstb.2018.0547

Koštál, V. (2006). Eco-physiological phases of insect diapause. Journal of Insect Physiology, 52(2), 113–127. https://doi.org/10.1016/j.jinsphys.2005.09.008

Kozak, G. M., Wadsworth, C. B., Kahne, S. C., Bogdanowicz, S. M., Harrison, R. G., Coates, B. S., & Dopman, E. B. (2019). Genomic Basis of Circannual Rhythm in the European Corn Borer Moth. Current Biology, 29(20), 3501–3509.e5. https://doi.org/10.1016/j.cub.2019.08.053

Kumar, S., Chen, D., Jang, C., Nall, A., Zheng, X., & Sehgal, A. (2014). An ecdysone-responsive nuclear receptor regulates circadian rhythms in Drosophila. Nature Communications, 5, 5697. https://doi.org/10.1038/ncomms6697

Kyriacou, C. P., Peixoto, A. A., Sandrelli, F., Costa, R., & Tauber, E. (2008). Clines in clock genes: Fine-tuning circadian rhythms to the environment. Trends in Genetics, 24(3), 124–132. https://doi.org/10.1016/j.tig.2007.12.003

Lachapelle, J., Reid, J., & Colegrave, N. (2015). Repeatability of adaptation in experimental populations of different sizes. Proceedings of the Royal Society B: Biological Sciences, 282(1805), 20143033. https://doi.org/10.1098/rspb.2014.3033

Langmead, B., & Salzberg, S. L. (2012). Fast gapped-read alignment with Bowtie 2. Nature Methods, 9(4), 357–359. https://doi.org/10.1038/nmeth.1923

Larkin, A., Marygold, S. J., Antonazzo, G., Attrill, H., dos Santos, G., Garapati, P. V., Goodman, J. L., Gramates, L. S., Millburn, G., Strelets, V. B., Tabone, C. J., Thurmond, J., FlyBase Consortium, Perrimon, N., Gelbart, S. R., Agapite, J., Broll, K., Crosby, M., dos Santos, G., … Lovato, T. (2021). FlyBase: Updates to the Drosophila melanogaster knowledge base. Nucleic Acids Research, 49(D1), D899–D907. https://doi.org/10.1093/nar/gkaa1026

Lee, S. F., Sgrò, C. M., Shirriffs, J., Wee, C. W., Rako, L., Van HEERWAARDEN, B., & Hoffmann, A. A. (2011). Polymorphism in the couch potato gene clines in eastern Australia but is not associated with ovarian dormancy in Drosophila melanogaster. Molecular Ecology, 20(14), 2973–2984. https://doi.org/10.1111/j.1365-294X.2011.05155.x

Levy, R. C., Kozak, G. M., Wadsworth, C. B., Coates, B. S., & Dopman, E. B. (2015). Explaining the sawtooth: Latitudinal periodicity in a circadian gene correlates with shifts in generation number. Journal of Evolutionary Biology, 28(1), 40–53. https://doi.org/10.1111/jeb.12562

Li, H., Handsaker, B., Wysoker, A., Fennell, T., Ruan, J., Homer, N., Marth, G., Abecasis, G., Durbin, R., & 1000 Genome Project Data Processing Subgroup. (2009). The Sequence Alignment/Map format and SAMtools. Bioinformatics (Oxford, England), 25(16), 2078–2079. https://doi.org/10.1093/bioinformatics/btp352

Li, K., Tian, L., Guo, Z., Guo, S., Zhang, J., Gu, S.-H., Palli, S. R., Cao, Y., & Li, S. (2016). 20-Hydroxyecdysone (20E) Primary Response Gene E75 Isoforms Mediate Steroidogenesis Autoregulation and Regulate Developmental Timing in Bombyx*. Journal of Biological Chemistry, 291(35), 18163–18175. https://doi.org/10.1074/jbc.M116.737072

Li, N., Tong, X., Zeng, J., Meng, G., Sun, F., Hu, H., Song, J., Lu, C., & Dai, F. (2019). Hippo pathway regulates somatic development and cell proliferation of silkworm. Genomics, 111(3), 391–397. https://doi.org/10.1016/j.ygeno.2018.02.014

Li, W.-D., Chen, S.-X., & Qin, J.-G. (2003). Yazhou yumiming yu Ouzhou yumiming hunshengqu de yanjiu [Identification of sympatric Asian and European corn borers]. Entomological Knowledge, 40(1), 31–35.

Lopez, L., Fasano, C., Perrella, G., & Facella, P. (2021). Cryptochromes and the Circadian Clock: The Story of a Very Complex Relationship in a Spinning World. Genes, 12(5), 672. https://doi.org/10.3390/genes12050672

Ma, R., Qian, H. T., Dong, H., Xia, X., & Cong, B. (2008). Butong dili zhongqun Yazhou yumiming yuedong youchong fusuhou de fayu liqi yanjiu [Research on the Development Period of Over-wintering Larvae of Different Geographic Populations of Aisan Corn Borer]. Hubei Agricultural Sciences, 47(5), 541–543.

Machado, H. E., Bergland, A. O., O’brien, K. R., Behrman, E. L., Schmidt, P. S., & Petrov, D. A. (2016). Comparative population genomics of latitudinal variation in Drosophila simulans and Drosophila melanogaster. Molecular Ecology, 25(3), 723. https://doi.org/10.1111/mec.13446

Mallet, J. (2005). Hybridization as an invasion of the genome. Trends in Ecology & Evolution, 20(5), 229–237. https://doi.org/10.1016/j.tree.2005.02.010

Martin, S. H. (2018). GitHub—Simonhmartin/genomics_general: General tools for genomic analyses. GitHub. https://github.com/simonhmartin/genomics_general

Martin, S. H., Davey, J. W., & Jiggins, C. D. (2015). Evaluating the Use of ABBA–BABA Statistics to Locate Introgressed Loci. Molecular Biology and Evolution, 32(1), 244–257. https://doi.org/10.1093/molbev/msu269

Mathias, D., Jacky, L., Bradshaw, W. E., & Holzapfel, C. M. (2005). Geographic and developmental variation in expression of the circadian rhythm gene, timeless, in the pitcher-plant mosquito, Wyeomyia smithii. Journal of Insect Physiology, 51(6), 661–667. https://doi.org/10.1016/j.jinsphys.2005.03.011

McLeod, D. G. R. (1976). Geographical variation of diapause termination in the European corn borer, Ostrinia nubilalis (Lepidoptera: Pyralidae), in southwestern Ontario. The Canadian Entomologist, 108(12), 1403– 1408. https://doi.org/10.4039/Ent1081403-12

Mi, H., Muruganujan, A., Huang, X., Ebert, D., Mills, C., Guo, X., & Thomas, P. D. (2019). Protocol Update for large-scale genome and gene function analysis with the PANTHER classification system (v.14.0). *Nature Protocols*, *14*(3), 703–721. https://doi.org/10.1038/s41596-019-0128-8

Michel, A. P., Rull, J., Aluja, M., & Feder, J. L. (2007). The genetic structure of hawthorn-infesting Rhagoletis pomonella populations in Mexico: Implications for sympatric host race formation. Molecular Ecology, 16(14), 2867–2878. https://doi.org/10.1111/j.1365-294X.2007.03263.x

Murillo-Maldonado, J. M., Zeineddine, F. B., Stock, R., Thackeray, J., & Riesgo-Escovar, J. R. (2011). Insulin Receptor-Mediated Signaling via Phospholipase C-γ Regulates Growth and Differentiation in Drosophila. PLOS ONE, 6(11), e28067. https://doi.org/10.1371/journal.pone.0028067

Nachman, M. W., Hoekstra, H. E., & D’Agostino, S. L. (2003). The genetic basis of adaptive melanism in pocket mice. Proceedings of the National Academy of Sciences, 100(9), 5268–5273. https://doi.org/10.1073/pnas.0431157100

Nagy, D., Andreatta, G., Bastianello, S., Martín Anduaga, A., Mazzotta, G., Kyriacou, C. P., & Costa, R. (2018). A Semi-natural Approach for Studying Seasonal Diapause in Drosophila melanogaster Reveals Robust Photoperiodicity. Journal of Biological Rhythms, 33(2), 117–125. https://doi.org/10.1177/0748730417754116

Oh, H., & Irvine, K. D. (2010). Yorkie: The final destination of Hippo signaling. Trends in Cell Biology, 20(7), 410–417. https://doi.org/10.1016/j.tcb.2010.04.005

Pan, X., & O’Connor, M. B. (2021). Coordination among multiple receptor tyrosine kinase signals controls Drosophila developmental timing and body size. Cell Reports, 36(9), 109644. https://doi.org/10.1016/j.celrep.2021.109644

Paolucci, S., Salis, L., Vermeulen, C. J., Beukeboom, L. W., & van de Zande, L. (2016). QTL analysis of the photoperiodic response and clinal distribution of period alleles in Nasonia vitripennis. Molecular Ecology, 25(19), 4805–4817. https://doi.org/10.1111/mec.13802

Paolucci, S., van de Zande, L., & Beukeboom, L. W. (2013). Adaptive latitudinal cline of photoperiodic diapause induction in the parasitoid Nasonia vitripennis in Europe. Journal of Evolutionary Biology, 26(4), 705–718. https://doi.org/10.1111/jeb.12113

Paradis, E., & Schliep, K. (2019). ape 5.0: An environment for modern phylogenetics and evolutionary analyses in R. Bioinformatics, 35, 526–528.

Pegoraro, M., Zonato, V., Tyler, E. R., Fedele, G., Kyriacou, C. P., & Tauber, E. (2017). Geographical analysis of diapause inducibility in European Drosophila melanogaster populations. Journal of Insect Physiology, 98, 238–244. https://doi.org/10.1016/j.jinsphys.2017.01.015

Peypelut, L., Beydon, P., & Lavenseau, L. (1990). 20-hydroxyecdysone triggers the resumption of imaginal wing disc development after diapause in the European corn borer, Ostrinia nubilalis. Archives of Insect Biochemistry and Physiology, 15(1), 1–19. https://doi.org/10.1002/arch.940150102

Pfeifer, B., & Kapan, D. D. (2019). Estimates of introgression as a function of pairwise distances. BMC Bioinformatics, 20(1), 207. https://doi.org/10.1186/s12859-019-2747-z

Picard Toolkit. (2019). Broad Institute; GitHub Repository. http://broadinstitute.github.io/picard/

Posledovich, D., Toftegaard, T., Navarro-Cano, J. A., Wiklund, C., Ehrlén, J., & Gotthard, K. (2014). Latitudinal variation in thermal reaction norms of post-winter pupal development in two butterflies differing in phenological specialization. Biological Journal of the Linnean Society, 113(4), 981–991. https://doi.org/10.1111/bij.12371

Pruisscher, P., Nylin, S., Gotthard, K., & Wheat, C. W. (2018). Genetic variation underlying local adaptation of diapause induction along a cline in a butterfly. Molecular Ecology, 27(18), 3613–3626. https://doi.org/10.1111/mec.14829

Quinlan, A. R., & Hall, I. M. (2010). BEDTools: A flexible suite of utilities for comparing genomic features. Bioinformatics, 26(6), 841–842. https://doi.org/10.1093/bioinformatics/btq033

R Core Team. (2020). R: A language and environment for statistical computing (4.0.0) [Computer software]. R Foundation for Statistical Computing. https://www.R-project.org/

Ragland, G. J., Armbruster, P. A., & Meuti, M. E. (2019). Evolutionary and functional genetics of insect diapause: A call for greater integration. Current Opinion in Insect Science, 36, 74–81. https://doi.org/10.1016/j.cois.2019.08.003

Ragland, G. J., Doellman, M. M., Meyers, P. J., Hood, G. R., Egan, S. P., Powell, T. H. Q., Hahn, D. A., Nosil, P., & Feder, J. L. (2017). A test of genomic modularity among life-history adaptations promoting speciation with gene flow. Molecular Ecology, 26(15), 3926–3942. https://doi.org/10.1111/mec.14178

Raudvere, U., Kolberg, L., Kuzmin, I., Arak, T., Adler, P., Peterson, H., & Vilo, J. (2019). g:Profiler: A web server for functional enrichment analysis and conversions of gene lists (2019 update). Nucleic Acids Research, 47(W1), W191–W198. https://doi.org/10.1093/nar/gkz369

Regier, J. C., Fang, Q. Q., Mitter, C., Peigler, R. S., Friedlander, T. P., & Solis, M. A. (1998). Evolution and phylogenetic utility of the period gene in Lepidoptera. Molecular Biology and Evolution, 15(9), 1172–1182.

Reinhardt, J. A., Kolaczkowski, B., Jones, C. D., Begun, D. J., & Kern, A. D. (2014). Parallel Geographic Variation in Drosophila melanogaster. Genetics, 197(1), 361–373. https://doi.org/10.1534/genetics.114.161463

Rennison, D. J., & Peichel, C. L. (2022). Pleiotropy facilitates parallel adaptation in sticklebacks. Molecular Ecology, 31(5), 1476–1486. https://doi.org/10.1111/mec.16335

Riddiford, L. M. (1993). Hormone receptors and the regulation of insect metamorphosis. Receptor, 3(3), 203–209.

Roff, D. (1980). Optimizing development time in a seasonal environment: The ‘ups and downs’ of clinal variation. Oecologia, 45(2), 202–208. https://doi.org/10.1007/BF00346461

Rousset, F. (1997). Genetic Differentiation and Estimation of Gene Flow from FStatistics Under Isolation by Distance. Genetics, 145, 1219–1228.

Sadakiyo, S., & Ishihara, M. (2011). Rapid seasonal adaptation of an alien bruchid after introduction: Geographic variation in life cycle synchronization and critical photoperiod for diapause induction. Entomologia Experimentalis et Applicata, 140(1), 69–76. https://doi.org/10.1111/j.1570-7458.2011.01136.x

Saez, L., Derasmo, M., Meyer, P., Stieglitz, J., & Young, M. W. (2011). A Key Temporal Delay in the Circadian Cycle of Drosophila Is Mediated by a Nuclear Localization Signal in the Timeless Protein. Genetics, 188(3), 591–600. https://doi.org/10.1534/genetics.111.127225

Sandrelli, F., Tauber, E., Pegoraro, M., Mazzotta, G., Cisotto, P., Landskron, J., Stanewsky, R., Piccin, A., Rosato, E., Zordan, M., Costa, R., & Kyriacou, C. P. (2007). A Molecular Basis for Natural Selection at the timeless Locus in Drosophila melanogaster. Science, 316(5833), 1898–1900. https://doi.org/10.1126/science.1138426

Sathyanarayanan, S., Zheng, X., Xiao, R., & Sehgal, A. (2004). Posttranslational Regulation of Drosophila PERIOD Protein by Protein Phosphatase 2A. Cell, 116(4), 603–615. https://doi.org/10.1016/S0092-8674(04)00128-X

Saunders, D. S. (2010). Controversial aspects of photoperiodism in insects and mites. Journal of Insect Physiology, 56(11), 1491–1502. https://doi.org/10.1016/j.jinsphys.2010.05.002

Saunders, D. S. (2012). Insect photoperiodism: Seeing the light. Physiological Entomology, 37(3), 207–218. https://doi.org/10.1111/j.1365-3032.2012.00837.x

Saunders, D. S. (2016). The temporal ‘structure’ and function of the insect photoperiodic clock: A tribute to Colin S. Pittendrigh. Physiological Entomology, 41(1), 1–18. https://doi.org/10.1111/phen.12131

Saunders, D. S. (2020). Dormancy, Diapause, and the Role of the Circadian System in Insect Photoperiodism. Annual Review of Entomology, 65(1), 373–389. https://doi.org/10.1146/annurev-ento-011019-025116

Sawa, M., Kay, S. A., & Imaizumi, T. (2008). Photoperiodic flowering occurs under internal and external coincidence. Plant Signaling & Behavior, 3(4), 269–271.

Schlicker, A., Domingues, F. S., Rahnenführer, J., & Lengauer, T. (2006). A new measure for functional similarity of gene products based on Gene Ontology. BMC Bioinformatics, 7(1), 302. https://doi.org/10.1186/1471-2105-7-302

Schlötterer, C., Tobler, R., Kofler, R., & Nolte, V. (2014). Sequencing pools of individuals —Mining genome-wide polymorphism data without big funding. Nature Reviews Genetics, 15(11), 749–763. https://doi.org/10.1038/nrg3803

Schmidt, P. S., Matzkin, L., Ippolito, M., & Eanes, W. F. (2005). Geographic Variation in Diapause Incidence, Life-History Traits, and Climatic Adaptation in Drosophila Melanogaster. Evolution, 59(8), 1721–1732. https://doi.org/10.1111/j.0014-3820.2005.tb01821.x

Schmidt, P. S., Zhu, C.-T., Das, J., Batavia, M., Yang, L., & Eanes, W. F. (2008). An amino acid polymorphism in the couch potato gene forms the basis for climatic adaptation in Drosophila melanogaster. Proceedings of the National Academy of Sciences, 105(42), 16207–16211. https://doi.org/10.1073/pnas.0805485105

Shama, L. N. S., Campero-Paz, M., Wegner, K. M., De Block, M., & Stoks, R. (2011). Latitudinal and voltinism compensation shape thermal reaction norms for growth rate. Molecular Ecology, 20(14), 2929–2941. https://doi.org/10.1111/j.1365-294X.2011.05156.x

Shimizu, T., & Kawasaki, K. (2001). Geographic variability in diapause response of Japanese Orius species. Entomologia Experimentalis et Applicata, 98(3), 303–316. https://doi.org/10.1046/j.1570-7458.2001.00787.x

Showers, W. B., Chiang, H. C., Keaster, A. J., Hill, R. E., Reed, G. L., Sparks, A. N., & Musick, G. J. (1975). Ecotypes of the European Corn Borer in North America. Environmental Entomology, 4(5), 753–760. https://doi.org/10.1093/ee/4.5.753

Sim, C., & Denlinger, D. (2013). Insulin signaling and the regulation of insect diapause. Frontiers in Physiology, 4. https://www.frontiersin.org/article/10.3389/fphys.2013.00189

Skopik, S. D., & Bowen, M. F. (1976). Insect photoperiodism: An hourglass measures photoperiodic time in Ostrinia nubilalis. Journal of Comparative Physiology, 111(3), 249–259. https://doi.org/10.1007/BF00606467

Slowikowski, K. (2021). *ggrepel: Automatically Position Non-Overlapping Text Labels with “ggplot2”* (R package version 0.9.1) [Computer software]. https://CRAN.R-project.org/package=ggrepel

Smith, D. C. (1988). Heritable divergence of Rhagoletis pomonella host races by seasonal asynchrony. Nature, 336(6194), 66–67. https://doi.org/10.1038/336066a0

Smith, J., & Kronforst, M. R. (2013). Do Heliconius butterfly species exchange mimicry alleles? Biology Letters, 9(4), 20130503. https://doi.org/10.1098/rsbl.2013.0503

Song, Y. H., Ito, S., & Imaizumi, T. (2010). Similarities in the circadian clock and photoperiodism in plants. Current Opinion in Plant Biology, 13(5), 594–603. https://doi.org/10.1016/j.pbi.2010.05.004

Sota, T. (1994). Larval diapause, size, and autogeny in the mosquito Aedes togoi (Diptera, Culicidae) from tropical to subarctic zones. Canadian Journal of Zoology, 72(8), 1462–1468. https://doi.org/10.1139/z94-193

Sourvorov, A., Kapustin, Y., Kiryutin, B., Chetvernin, V., Tatusova, T., & Lipman, D. (2010, February 25). *Gnomon—The NCBI eukaryotic gene prediction tool [online]*. National Center for Biotechnology Information. https://www.ncbi.nlm.nih.gov/core/assets/genome/files/Gnomon-description.pdf

Sparks, A. N., & Young, J. R. (1971). European corn borer activity in Georgia. Journal of the Georgia Entomological Society, 6, 211–215.

Steiner, C. C., Weber, J. N., & Hoekstra, H. E. (2007). Adaptive Variation in Beach Mice Produced by Two Interacting Pigmentation Genes. PLOS Biology, 5(9), e219. https://doi.org/10.1371/journal.pbio.0050219

Stern, D. L., & Orgogozo, V. (2009). Is Genetic Evolution Predictable? Science, 323(5915), 746–751. https://doi.org/10.1126/science.1158997

Storz, J. F. (2016). Causes of molecular convergence and parallelism in protein evolution. Nature Reviews Genetics, 17(4), 239–250. https://doi.org/10.1038/nrg.2016.11

Supek, F., Bošnjak, M., Škunca, N., & Šmuc, T. (2011). REVIGO Summarizes and Visualizes Long Lists of Gene Ontology Terms. PLOS ONE, 6(7), e21800. https://doi.org/10.1371/journal.pone.0021800

Sutherland, W. J., Freckleton, R. P., Godfray, H. C. J., Beissinger, S. R., Benton, T., Cameron, D. D., Carmel, Y., Coomes, D. A., Coulson, T., Emmerson, M. C., Hails, R. S., Hays, G. C., Hodgson, D. J., Hutchings, M. J., Johnson, D., Jones, J. P. G., Keeling, M. J., Kokko, H., Kunin, W. E., … Wiegand, T. (2013). Identification of 100 fundamental ecological questions. Journal of Ecology, 101(1), 58–67. https://doi.org/10.1111/1365-2745.12025

Takada, S., Akter, A., Itabashi, E., Nishida, N., Shea, D. J., Miyaji, N., Mehraj, H., Osabe, K., Shimizu, M., Takasaki-Yasuda, T., Kakizaki, T., Okazaki, K., Dennis, E. S., & Fujimoto, R. (2019). The role of FRIGIDA and FLOWERING LOCUS C genes in flowering time of Brassica rapa leafy vegetables. Scientific Reports, 9(1), 13843. https://doi.org/10.1038/s41598-019-50122-2

Takeda, M., & Skopik, S. D. (1985). Geographic variation in the circadian system controlling photoperiodism in Ostrinia nubilalis. Journal of Comparative Physiology A, 156(5), 653–658. https://doi.org/10.1007/BF00619114

Tauber, E., Zordan, M., Sandrelli, F., Pegoraro, M., Osterwalder, N., Breda, C., Daga, A., Selmin, A., Monger, K., Benna, C., Rosato, E., Kyriacou, C. P., & Costa, R. (2007). Natural selection favors a newly derived timeless allele in Drosophila melanogaster. Science (New York, N.Y.), 316(5833), 1895–1898. https://doi.org/10.1126/science.1138412

Tauber, M. J., Tauber, C. A., & Masaki, S. (1986). Seasonal Adaptations of Insects. Oxford University Press.

Tenaillon, M. I., & Charcosset, A. (2011). A European perspective on maize history. Comptes Rendus Biologies, 334(3), 221–228. https://doi.org/10.1016/j.crvi.2010.12.015

Thackeray, J. R., Gaines, P. C., Ebert, P., & Carlson, J. R. (1998). Small wing encodes a phospholipase C-(gamma) that acts as a negative regulator of R7 development in Drosophila. Development (Cambridge, England), 125(24), 5033–5042. https://doi.org/10.1242/dev.125.24.5033

Timer, J., Tobin, P. C., & Saunders, M. C. (2010). Geographic variation in diapause induction: The grape berry moth (Lepidoptera: Tortricidae). https://doi.org/10.1603/EN10116

Urbanski, J., Mogi, M., O’Donnell, D., DeCotiis, M., Toma, T., & Armbruster, P. (2012). Rapid Adaptive Evolution of Photoperiodic Response during Invasion and Range Expansion across a Climatic Gradient. The American Naturalist, 179(4), 490–500. https://doi.org/10.1086/664709

Wadsworth, C. B., & Dopman, E. B. (2015). Transcriptome profiling reveals mechanisms for the evolution of insect seasonality. Journal of Experimental Biology, 218(22), 3611–3622. https://doi.org/10.1242/jeb.126136

Wadsworth, C. B., Okada, Y., & Dopman, E. B. (2020). Phenology-dependent cold exposure and thermal performance of Ostrinia nubilalis ecotypes. BMC Evolutionary Biology, 20(1), 34. https://doi.org/10.1186/s12862-020-1598-6

Wang, S., Lu, Y., Yin, M.-X., Wang, C., Wu, W., Li, J., Wu, W., Ge, L., Hu, L., Zhao, Y., & Zhang, L. (2016). Importin α1 Mediates Yorkie Nuclear Import via an N-terminal Non-canonical Nuclear Localization Signal*. Journal of Biological Chemistry, 291(15), 7926–7937. https://doi.org/10.1074/jbc.M115.700823

Wang, X.-P., Yang, Q.-S., Dalin, P., Zhou, X.-M., Luo, Z.-W., & Lei, C.-L. (2012). Geographic variation in photoperiodic diapause induction and diapause intensity in Sericinus montelus (Lepidoptera: Papilionidae). Insect Science, 19(3), 295–302. https://doi.org/10.1111/j.1744-7917.2011.01473.x

Wang, Y., Kim, K. S., Guo, W., Li, Q., Zhang, Y., Wang, Z., & Coates, B. S. (2017). Introgression between divergent corn borer species in a region of sympatry: Implications on the evolution and adaptation of pest arthropods. Molecular Ecology, 26(24), 6892–6907. https://doi.org/10.1111/mec.14387

Wickham, H. (2016). *ggplot2: Elegant Graphics for Data Analysis*. Springer-Verlag New York. https://ggplot2.tidyverse.org

Wickham, H. (2020). *tidyr: Tidy Messy Data* (R package version 1.1.2) [Computer software]. https://CRAN.R-project.org/package=tidyr

Wickham, H., François, R., Henry, L., & Müller, K. (2020). *dplyr: A Grammar of Data Manipulation* (R package version 1.0.2) [Computer software]. https://CRAN.R-project.org/package=dplyr

Wilke, C. O. (2020). *cowplot: Streamlined Plot Theme and Plot Annotations for “ggplot2”* (R package version 1.1.1) [Computer software]. https://CRAN.R-project.org/package=cowplot

Williams, C. M., Ragland, G. J., Betini, G., Buckley, L. B., Cheviron, Z. A., Donohue, K., Hereford, J., Humphries, M. M., Lisovski, S., Marshall, K. E., Schmidt, P. S., Sheldon, K. S., Varpe, Ø., & Visser, M. E. (2017). Understanding Evolutionary Impacts of Seasonality: An Introduction to the Symposium. Integrative and Comparative Biology, 57(5), 921–933. https://doi.org/10.1093/icb/icx122

Xiao, L., He, H.-M., Huang, L.-L., Geng, T., Fu, S., & Xue, F.-S. (2016). Variation of life-history traits of the Asian corn borer, Ostrinia furnacalis in relation to temperature and geographical latitude. Ecology and Evolution, 6(15), 5129–5143. https://doi.org/10.1002/ece3.2275

Yamahira, K., Kawajiri, M., Takeshi, K., & Irie, T. (2007). Inter- and Intrapopulation Variation in Thermal Reaction Norms for Growth Rate: Evolution of Latitudinal Compensation in Ectotherms with a Genetic Constraint. Evolution, 61(7), 1577–1589. https://doi.org/10.1111/j.1558-5646.2007.00130.x

Yang, H.-Z., Tu, X.-Y., Xia, Q.-W., He, H.-M., Chen, C., & Xue, F.-S. (2014). Photoperiodism of diapause induction and diapause termination in Ostrinia furnacalis. Entomologia Experimentalis et Applicata, 153(1), 34–46. https://doi.org/10.1111/eea.12226

Yasukochi, Y., Ohno, M., Shibata, F., Jouraku, A., Nakano, R., Ishikawa, Y., & Sahara, K. (2016). A FISH-based chromosome map for the European corn borer yields insights into ancient chromosomal fusions in the silkworm. Heredity, 116(1), 75–83. https://doi.org/10.1038/hdy.2015.72

Yoshii, T., Wülbeck, C., Sehadova, H., Veleri, S., Bichler, D., Stanewsky, R., & Helfrich -Förster, C. (2009). The Neuropeptide Pigment-Dispersing Factor Adjusts Period and Phase of Drosophila’s Clock. Journal of Neuroscience, 29(8), 2597–2610. https://doi.org/10.1523/JNEUROSCI.5439-08.2009

Yu, G. (2020). Using ggtree to visualize data on tree-like structures. Current Protocols in Bioinformatics, 69, e96. https://doi.org/10.1002/cpbi.96

Zonato, V., Collins, L., Pegoraro, M., Tauber, E., & Kyriacou, C. P. (2017). Is diapause an ancient adaptation in Drosophila? Journal of Insect Physiology, 98, 267–274. https://doi.org/10.1016/j.jinsphys.2017.01.017

