## Supplemental Table S1-S3, Figure S1-S2 for "Partial reuse of circadian clock genes along parallel clines of diapause in two moth species"

### Supplemental Methods

##### Calculation of genome-wide average *F*_ST_

For the genome-wide average *F*_ST_, the asymptotically consistent *F*_ST_ estimate (Bhatia et al., 2013) is the ratio of averages, not average of ratios, which means that the average of the single-SNP *F*_ST_ should not be used as the genome-wide average. However, due to limited computational resources, we could not calculate genome-wide average *F*_ST_ using the whole-genome data all at once in Popoolation2 (Kofler et al., 2011). Therefore, we divided the genome into 6 large windows (omitting the scaffold information) and calculated the average *F*_ST_ in each. The final genome-wide *F*_ST_ was the average of the 6 *F*_ST_ weighted by SNP number. We tried smaller windows and found that the weighted averages were similar to each other but were very different from the average of the single SNP *F*_ST_.

##### Correlation of covariate with population structure

For correlation between the covariates and the first Principal Component of the covariance matrix omega, we got the first PC of omega by singular value decomposition, following the code in BayPass’s R function simulate.PCcorrelated.covariate and Frachon et al., (2018). The specific R code used was cor.test(covariate, svd(omega)$u[ ,1], method=”Spearman”).

### Supplemental Tables and Figures

Table S1. Spearman correlation between covariates voltinism, critical day length (CDL), post-diapause development time (PDD), and between covariates and the population structure (first Principal Component of the covariance matrix omega). Lower triangle: Spearman’s rho. Upper triangle: Spearman’s p-value. Diagonal: Spearman’s rho between the covariate and the population structure.

|  | Latitude | Voltinism | CDL | PDD |
| --- | --- | --- | --- | --- |
| Latitude | **-0.07143** | 8.29E-05 | < 2.2E-16 | 0.3374 |
| Voltinism | -0.98198 | **0.03637** | 8.29E-05 | 0.3504 |
| CDL | 1 | -0.98198 | **-0.07143** | 0.3374 |
| PDD | 0.428571 | -0.41825 | 0.428571 | **-0.75** |

Table S2. SNPs showing evidence of adaptive introgression (in an f_d_ outlier window as well as associating with a phenological trait or being an XtX outlier).

| Scaffold | Position | Gene | Recipient population | Associated phenotype | XtX outlier |
| --- | --- | --- | --- | --- | --- |
| NW_021131247.1 | 19910 | *Got1* | HB_45N | Voltinism, CDL | No |
| NW_021135560.1 | 314603 | *Kap-alpha1* | HB_45N, SY_41N | Voltinism, PDD | Yes |
| NW_021135644.1 | 392159 | *ps* | HB_45N | PDD | No |
| NW_021132396.1 | 244964 | *Pdss2* | HF_31N | PDD | No |

Table S3. Alleles of select outlier SNPs in the clock gene per in ACB, ECB, Ostrinia scapulalis (ECB’s sibling species), and outgroup ALB. Outliers in ECB are as listed in Kozak et al. (2019) Supplementary Data S1. Outliers in ACB (this study) are selected based on whether they are XtX outliers, or, CDL/PDD-associated and nonsynonymous. O. scapulalis allele is based on one individual; ALB allele is based on three individuals. The second column gives the SNP’s position in the ACB scaffold **NW_021137771.1** and in the ECB scaffold **Scaffold532**, separated by “|”. Some positions are not covered in the outgroup, denoted by “-”. Alleles unique to ACB or ECB are bolded.

| Outlier SNP type | | Scaffold position  ACB \| ECB | ACB | ECB | *O. scapulalis* | ALB (ancestral) |
| --- | --- | --- | --- | --- | --- | --- |
| Kozak et al., (2019) | ECB: PLINK, E-box altering | 767507 \| 78628 | G | **C**/G | G | - |
|  | ECB: BayPass, in 5’ UTR | 745690 \| 93691 | G | **T**/G | - | G |
|  | ECB: PLINK, nonsynonymous | 689810 \| 140360 | C/T* | **A**/C | C | C/T |
|  | ECB: PLINK, nonsynonymous | 689801 \| 140369 | C/T* | C/T | C | T |
|  | ECB: PLINK, nonsynonymous | 689672 \| 140498 | A | A/**C** | A | A |
| This study | ACB: XtX, in intron | 705998 \| 124183 | **A**/C | C | G | - |
|  | ACB: XtX, nonsynon. (Arg to Trp) | 707355 \| 116467 | A/**T** | A | A | A |
|  | ACB: CDL, nonsynon. (Leu to Met) | 708691 \| 115084 | **A**/C | C | C | C |

*not significantly differentiated between populations in ACB

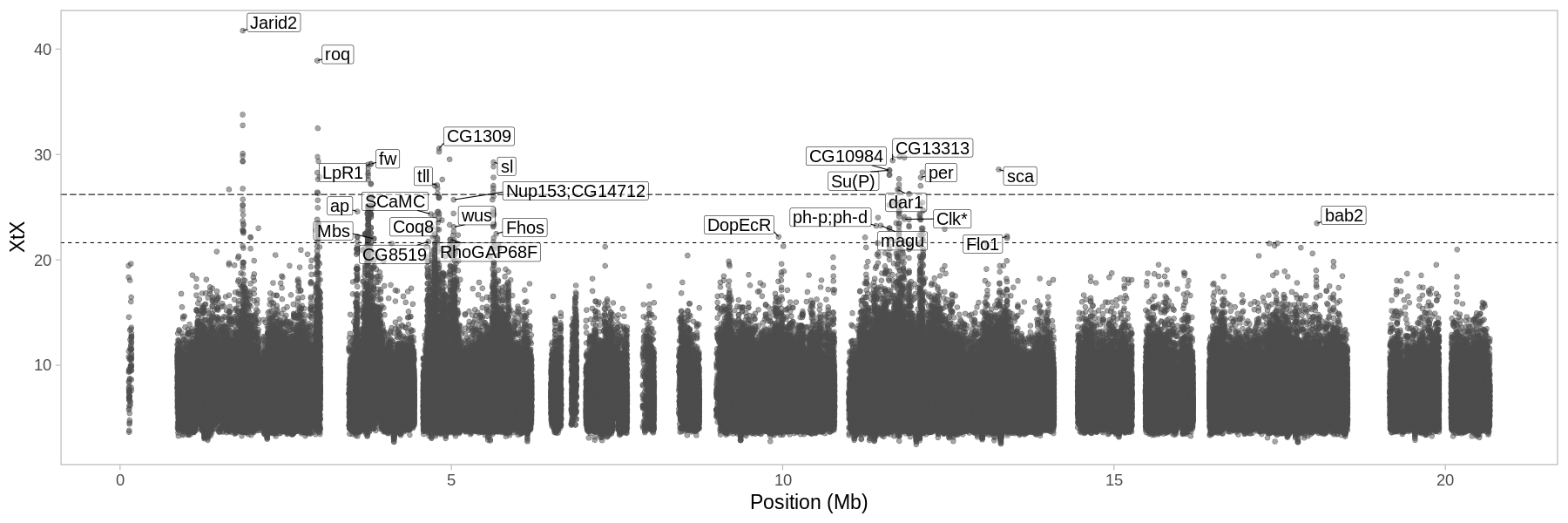

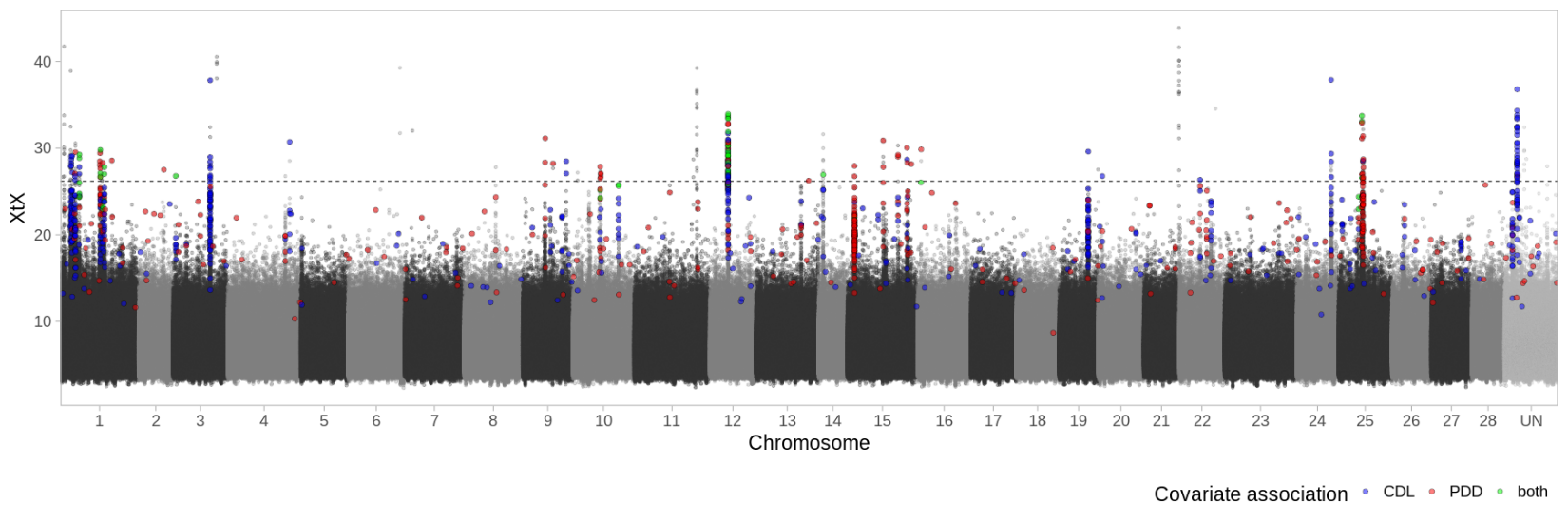

Figure S1. Genetic architecture of adaptation in ACB. Upper panel: XtX along the Z chromosome with the 99.99% and 99.9% thresholds (dashed lines). The 99.99% threshold was used for analyses in the main text. For other chromosomes see Supplemental File S2. Lower panel: XtX in all chromosomes with the 99.99% threshold as the dashed line. Additionally, SNPs significantly associated with CDL or PDD are colored. ACB scaffolds are aligned to the Bombyx mori chromosomes. UN: unplaced scaffolds. Clk*: predicted to be Clk (with predicted PAS domains) but there is another predicted Clk on Chromosome 4 (with a predicted bHLH-PAS_dClock domain); the two predicted proteins together align with the Bombyx mori CLK. Bab2: one of the two paralogs in Drosophila melanogaster (bab1 and bab2), but in Lepidopterans there is only one bab (Unbehend et al., 2021).

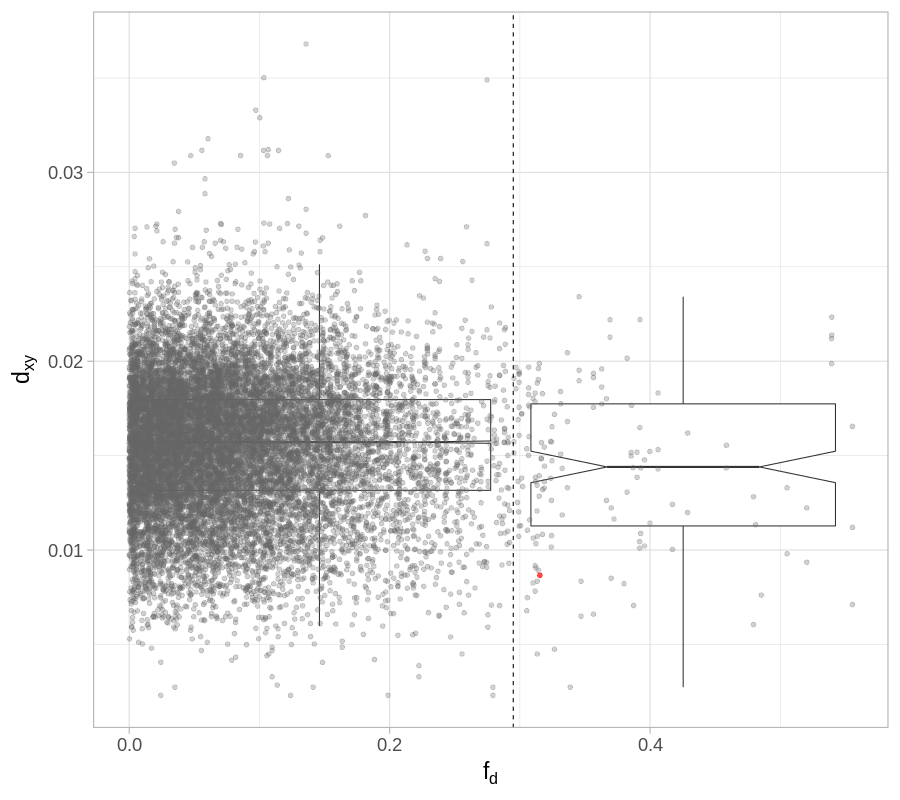

Figure S2. d_xy_ (HB_45N vs. ECB_PY) plotted against f_d_, with notched box plot comparing the distribution of d_xy_ in f_d_ outlier windows vs non-outlier windows. Dashed line: f_d_ outlier threshold (top 1%). Red: Kap-alpha1. d_xy_ was calculated in windows of 25,000 bp and f_d_ was calculated in windows of 300 SNPs.

### Supplemental References

Bhatia, G., Patterson, N., Sankararaman, S., & Price, A. L. (2013). Estimating and interpreting FST: The impact of rare variants. *Genome Research*, *23*(9), 1514–1521. https://doi.org/10.1101/gr.154831.113

Frachon, L., Bartoli, C., Carrère, S., Bouchez, O., Chaubet, A., Gautier, M., Roby, D., & Roux, F. (2018). A Genomic Map of Climate Adaptation in Arabidopsis thaliana at a Micro-Geographic Scale. *Frontiers in Plant Science*, *9*. https://doi.org/10.3389/fpls.2018.00967

Kofler, R., Pandey, R. V., & Schlötterer, C. (2011). PoPoolation2: Identifying differentiation between populations using sequencing of pooled DNA samples (Pool-Seq). *Bioinformatics*, *27*(24), 3435–3436. https://doi.org/10.1093/bioinformatics/btr589

Kozak, G. M., Wadsworth, C. B., Kahne, S. C., Bogdanowicz, S. M., Harrison, R. G., Coates, B. S., & Dopman, E. B. (2019). Genomic Basis of Circannual Rhythm in the European Corn Borer Moth. *Current Biology*, *29*(20), 3501-3509.e5. https://doi.org/10.1016/j.cub.2019.08.053

Unbehend, M., Kozak, G. M., Koutroumpa, F., Coates, B. S., Dekker, T., Groot, A. T., Heckel, D. G., & Dopman, E. B. (2021). Bric à brac controls sex pheromone choice by male European corn borer moths. *Nature Communications*, *12*(1), 2818. https://doi.org/10.1038/s41467-021-23026-x
