## Supplemental File S2 for "Partial reuse of circadian clock genes along parallel clines of diapause in two moth species"

**File S2.** XtX along all chromosomes (aligned to *Bombyx mori* genome, N=28) with the 99.99% and 99.9% thresholds (dashed lines) and genes labelled for SNPs above the 99.9% (lower) threshold. The 99.99% (higher) threshold was used for XtX outlier analyses in the main text. Chr 1 (the Z chromosome): see Figure S1 in File S1. The last row of plots here is the fused *B. mori* chromosomes (chr 11, 23, 24; see main text) as well as the unplaced scaffolds (in arbitrary order; no outlier genes with annotated functions hence no labels).

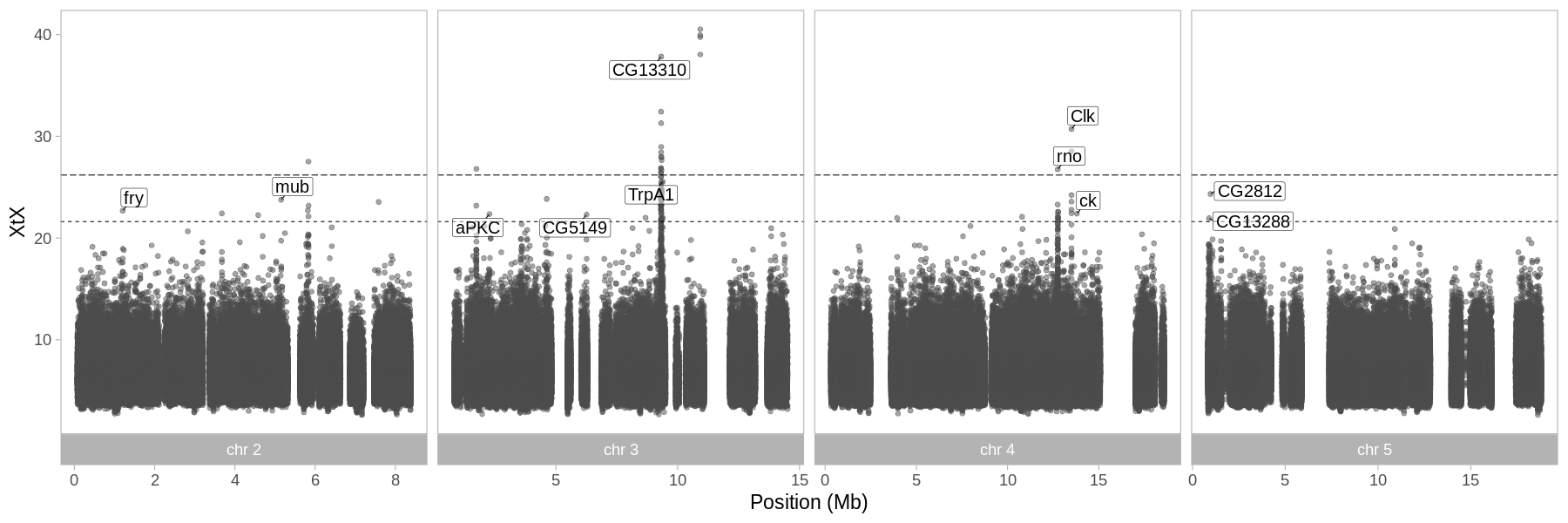

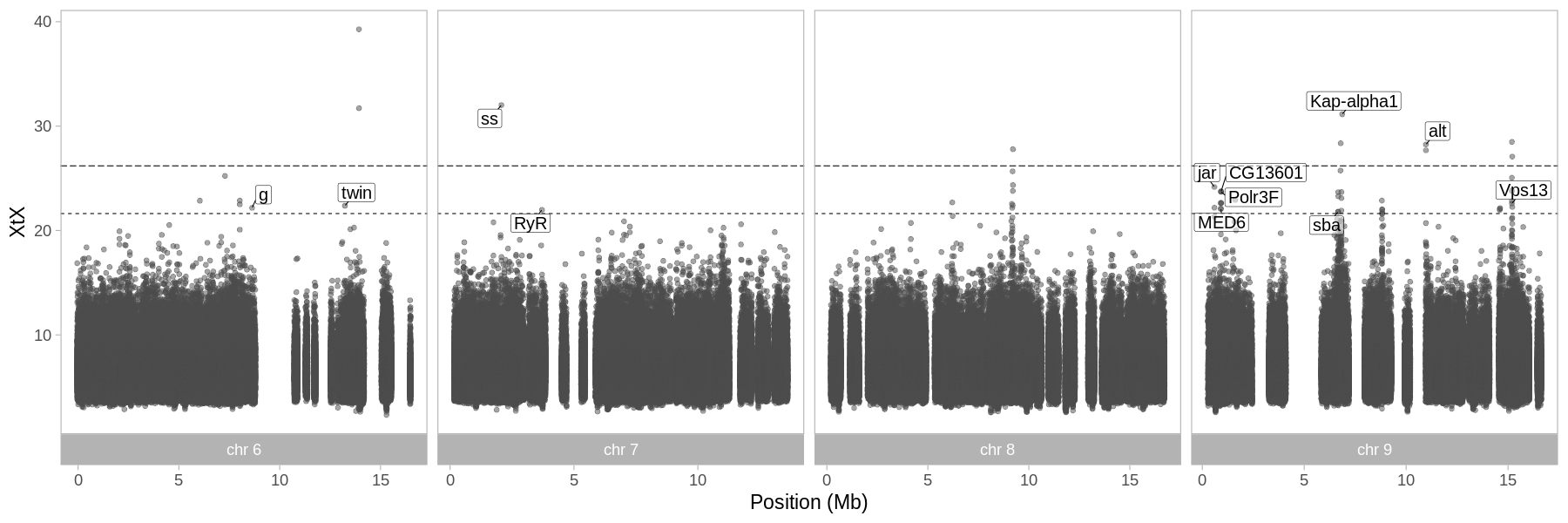

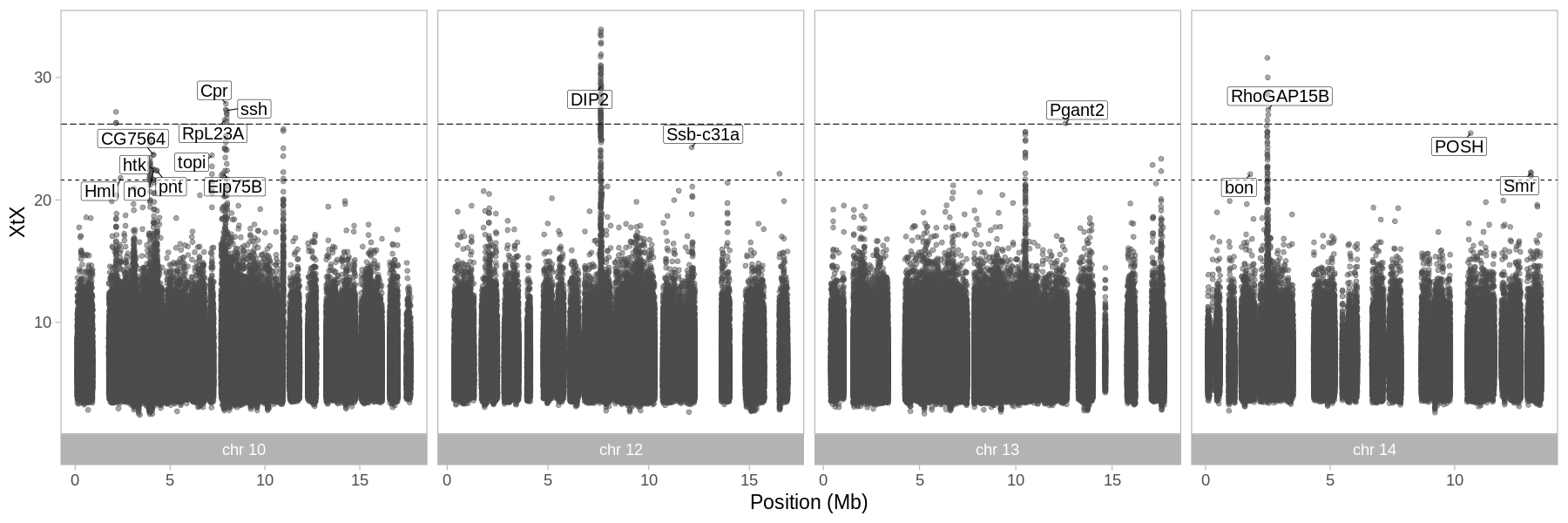

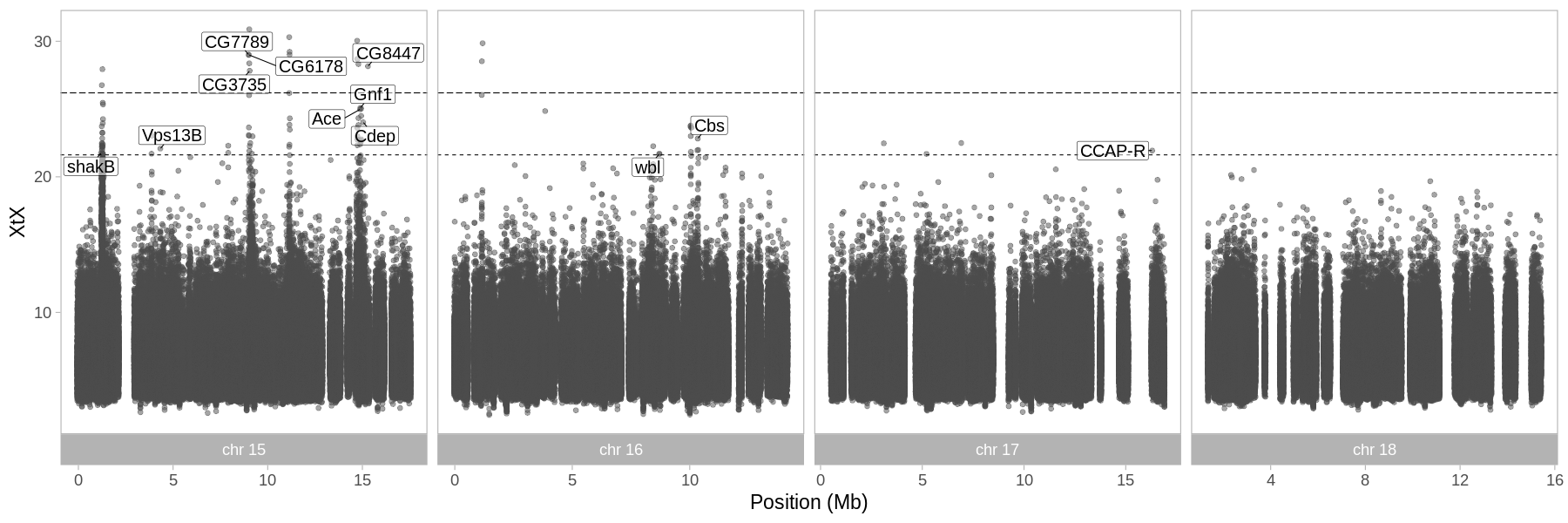

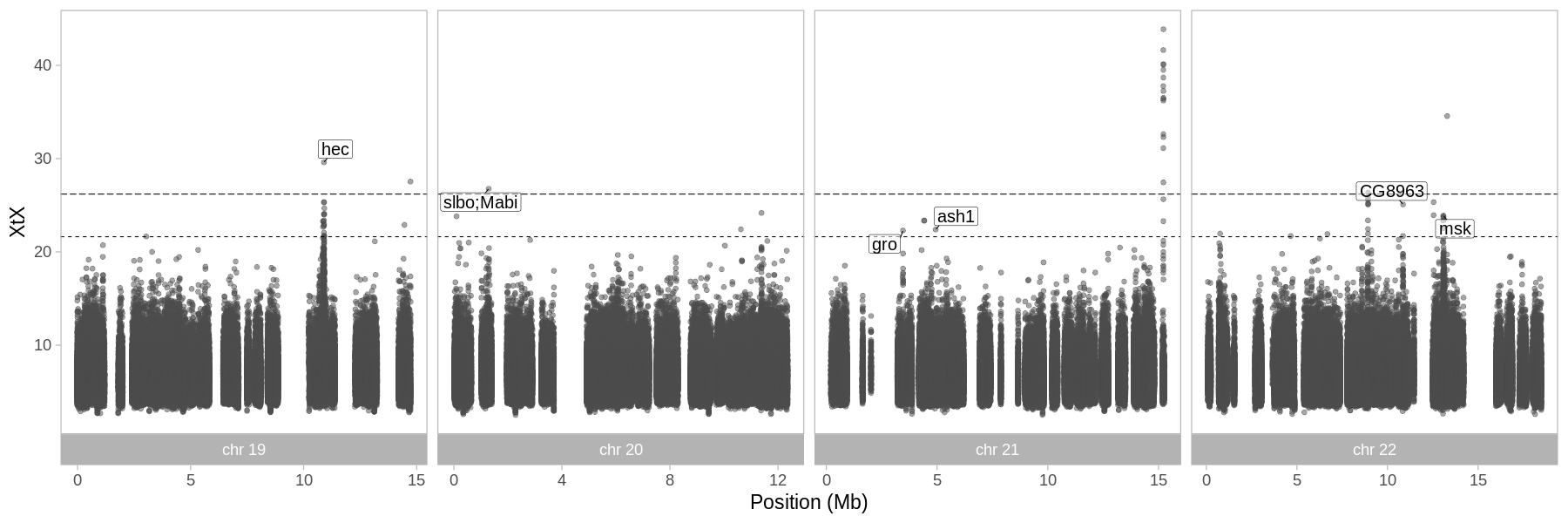

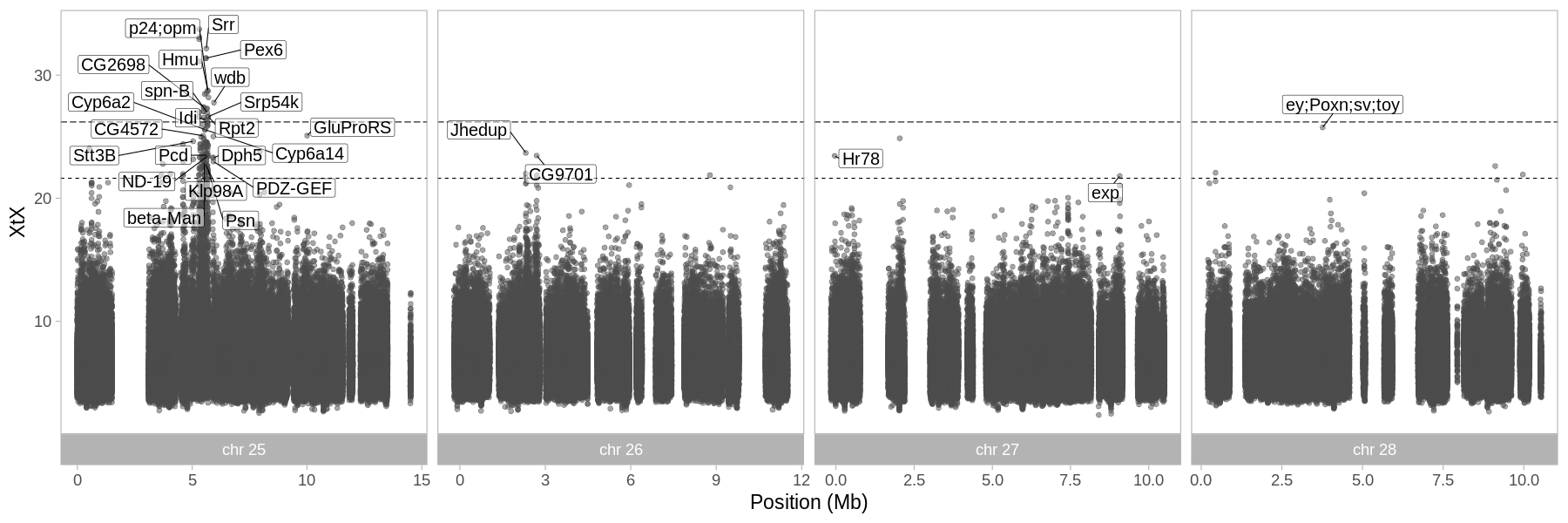

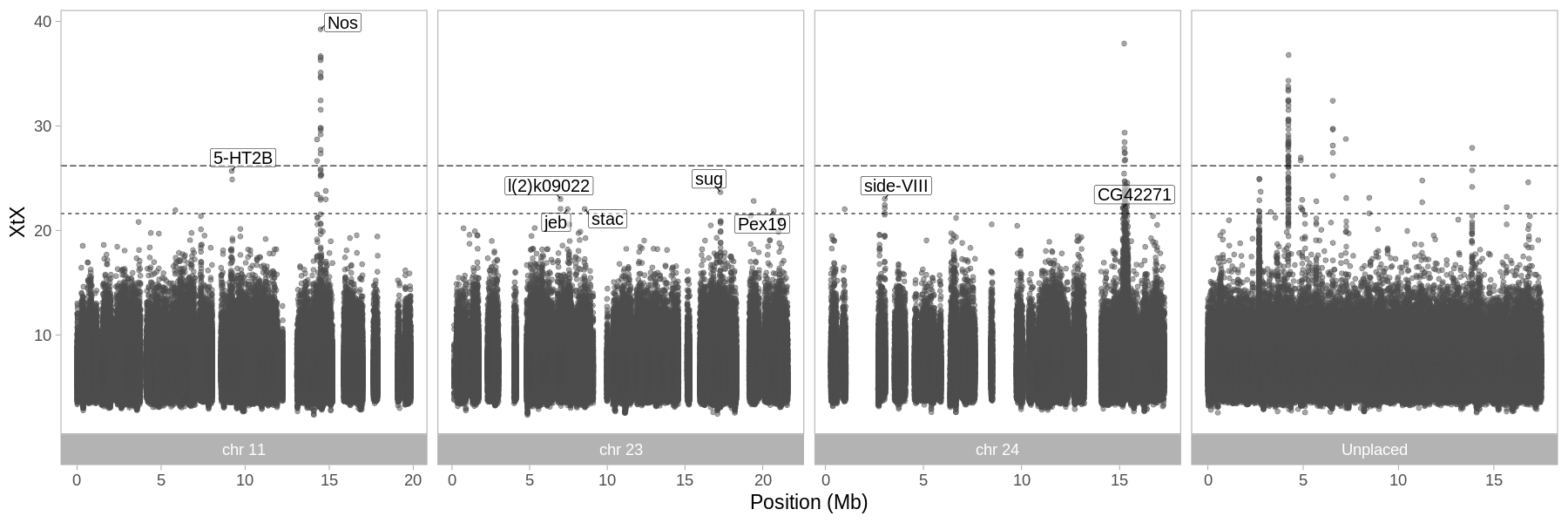
