## Supplemental File S4 for "Partial reuse of circadian clock genes along parallel clines of diapause in two moth species": gProfiler_sig_enriched_GOterms.docx

Comparison of all significantly enriched GO terms among the genes associated with critical daylength (CDL), post-diapause development time (PDD), voltinism, or latitude, or XtX outlier genes. Benjamin-Hochberg FDR-adjusted P values (g:Profiler; see main text) in strikethrough indicate that the GO term is not significantly enriched for that particular covariate/XtX.

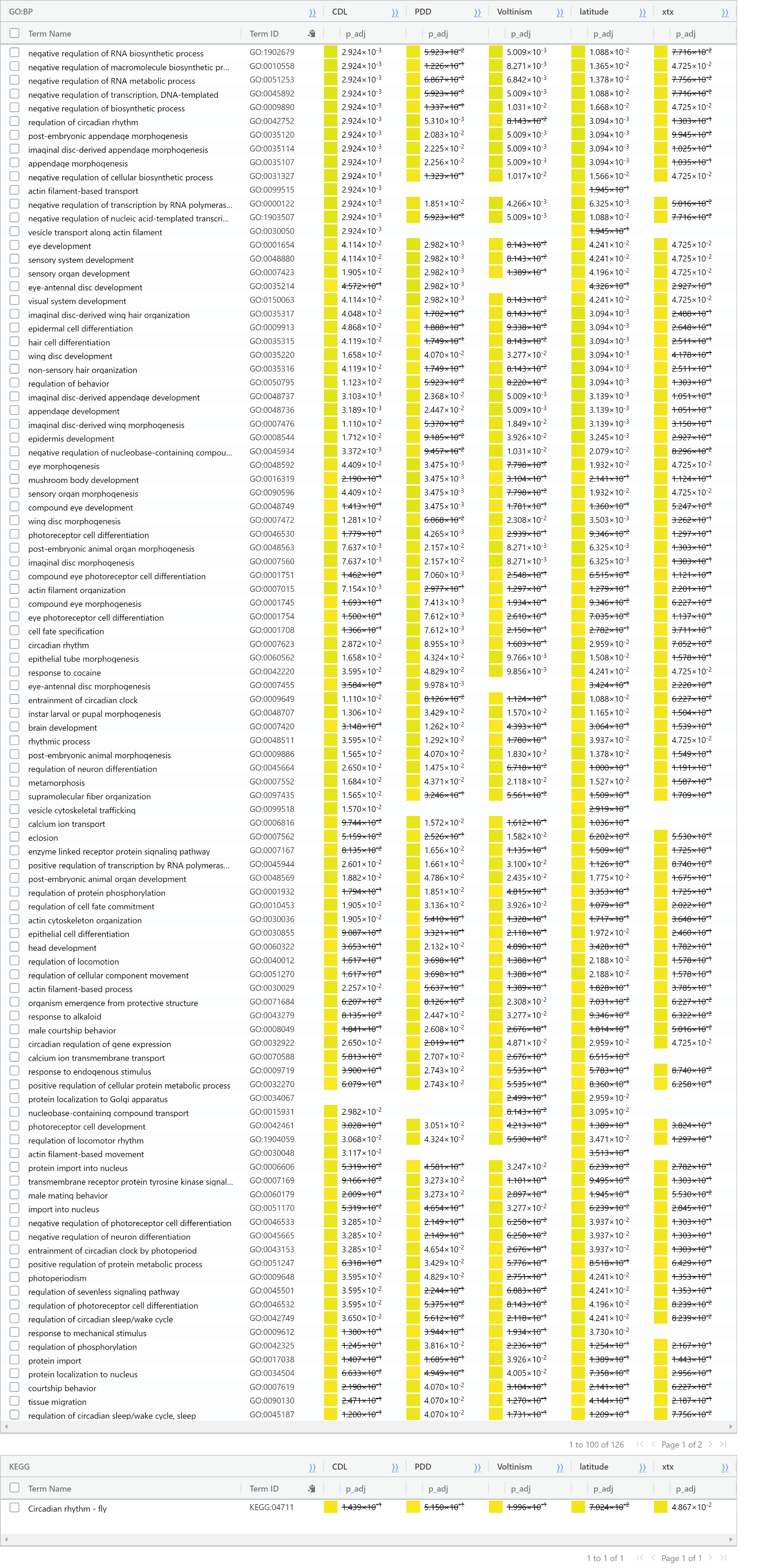

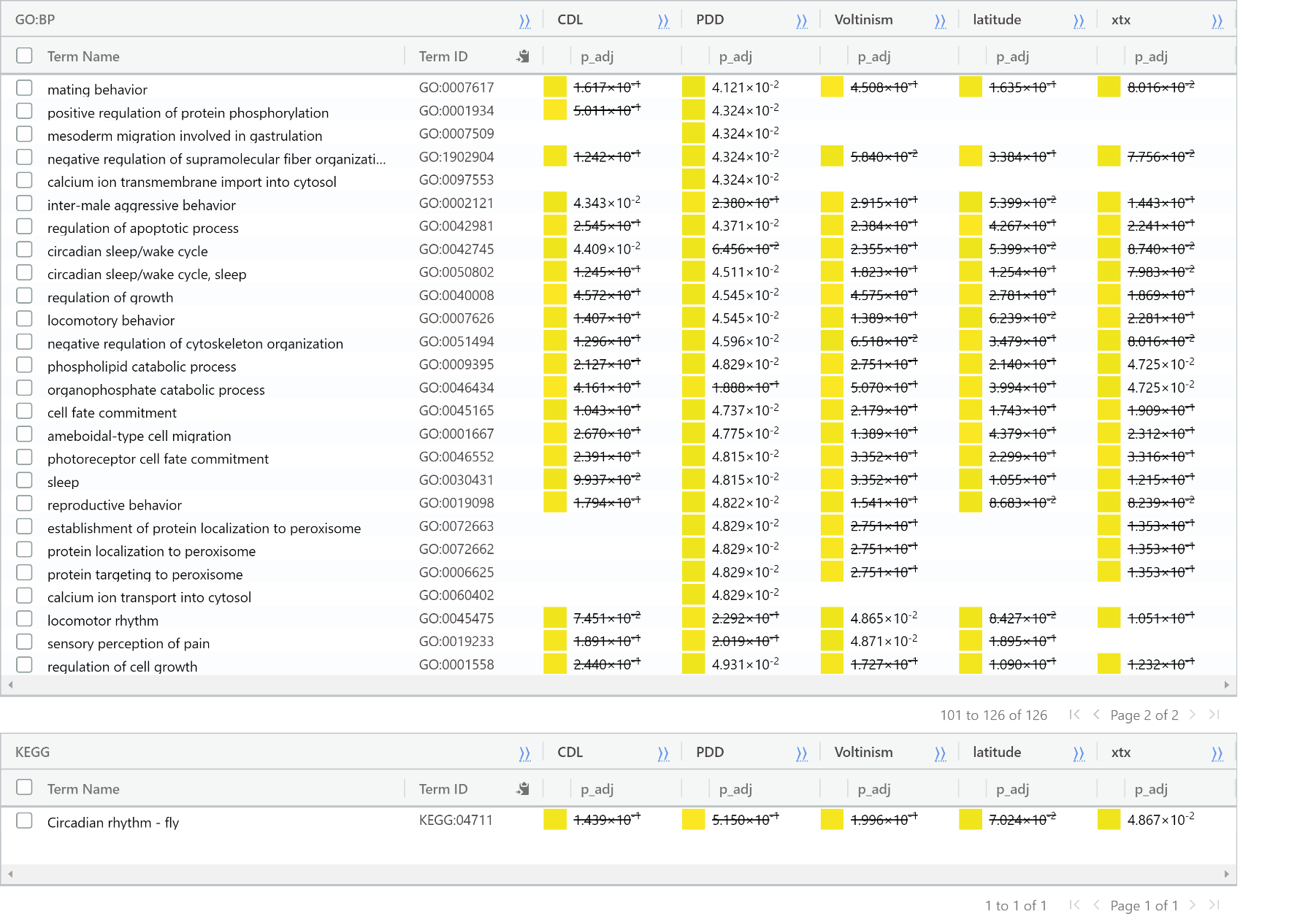
